# Identification, Characterization, and Targeting of a Rare and Temporal Dendritic Cell State that Facilitates Adaptive Immune Responses

**DOI:** 10.1101/2020.10.08.331744

**Authors:** Peter Deak, Bradley Studnitzer, Rachel Steinhardt, Aaron Esser-Kahn

## Abstract

The heterogeneity of innate immune cells facilitates efficient antigen presentation and immune activation in the presence of pathogens via cooperativity of various cell subsets and cell states but also obscures the contribution of individual antigen presenting cells (APCs) to overall immune response.^1^ It has been hypothesized that a small number of APCs, which are more sensitive to the initial pathogen stimulus, are responsible for coordinating neighboring APCs in an effort to share the metabolic strain associated with heightened pathogen sensitivity.^2^ In this study, we have identified a temporally-controlled state of dendritic cells (DCs) that demonstrate greater sensitivity to toll-like-receptor (TLR) agonists and secrete the majority of paracrine activating cytokines (TNFα, IL-6…ect). We were able to isolate this distinct population of DCs preferentially phagocytosed the majority of fluorescently labeled, TLR agonist conjugated microparticles (MPs).^3^ We call this population First Responder cells (FRs) due to their ability to first uptake the MPs and activate neighboring APCs via paracrine signaling. We show that FRs exist in this state for <3 hours, cycle through this state on a <24-hour timescale and show a distinct mRNA profile. Furthermore, FRs are necessary for generation of adaptive responses both in vitro and in vivo. We also show that we can improve both IgG titers and CD8 responses in vivo by targeting two highly upregulated receptors on FR cells, DAP12 and PRG2. Given the significance of FR involvement in APC activation, this study has broad immunological value because it offers a critical first evaluation of a new APC cell state but also has important translational value for improving vaccine efficacy via FR targeting.

## Introduction

Antigen presenting cells (APCs) initiate adaptive T and B cell response to pathogens by phagocytosing antigens and presenting antigen peptides on major histocompatibility type II molecules (MHCII). Given their importance for bridging the gap between adaptive and innate immunity, APCs are widely studied and are subdivided into various classical phenotypes of dendritic cells (DCs), B cells, Langerhans cells and macrophages.^4^ Of these, DCs are considered most critical for antigen presentation given their wide distribution in most dermal tissue, their migratory characteristics between dermal tissue and immune lymphoids and their high level of paracrine signaling which coordinates local immune responses.^5^ DCs are typically characterized by expression of CD11c and a high level of MHCII expression, however, DCs are also rather heterogenous and can be further divided into two main subtypes, a CD8+, XCR1+ subtype (cDC1) and a CD11b+, CD172+ subtype (cDC2).^6^ Given the diversity of DCs and subtypes, one question that continues to be elucidates is how these subtypes coordinate innate immune responses. Recent finding of sequencing, single-cell analysis, and microscopy have identified that within these population, there are distinct DCs states. Of particular interest to us was the identification that (<5%) of DC can, in certain conditions, express >70% of the inflammatory cytokine, TNFα, when simulated with a toll-like receptor (TLR) agonist, lipopolysaccharide (LPS).^2^ TLR signaling is an important co-stimulatory mechanism for facilitating antigen presentation in innate immune cells and promoting downstream adaptive responses to an antigen.^7^ These “super-secreting” cells appear to be one of the important cells for initiating global activation of a wider population of heterogeneous APCs. Study of these cells so far has been limited, due to lack of surface markers or methods of isolation.^8^

Inherent heterogeneity in immune cell populations contributing to a small subset of cells with wide ranging effects on broader populations is well established, as cell specialization and amplification of responses are traditionally hallmarks of immune cells. For example, CD4+ T cells are known to coordinate neighboring cell immunity via IL-2 secretion when stimulated with antigen.^9^ Likewise, innate cells have shown heterogeneity. Neutrophils, generally described as “sentinel cells” due to their active homing to pathogens, have been shown to be more heterogeneous and have subsets that seems to be more active in migration to pathogens.^10^ Even in a different subset of DCs, notably plasmacytoid DCs (pDC), there is increasing evidence that a small population of interferon (INF) secreting cells regulate bulk populations of pDC activation.^11^ Nevertheless, there is currently little direct evidence linking highly TLR reactive conventional DCs in a heterogeneous population and bulk activation of neighboring cells. In this article, we present evidence of a small population of APCs, which we refer to as First Responder DCs (FRs), which help facilitate bulk innate cell activation when cells are presented with limited TLR ligands. We identified and isolated these cells from heterogenous populations using fluorescently labeled TLR agonist conjugated microparticles (MPs). We observed that a small percentage (set at 5%) accumulated a statistically unlikely percentage (>90%) of TLR bearing particles when the ratio of particles to cells was 1:1. As these cells were highly fluorescent, we isolated FRs, observed unique transcriptional profiles, identified surface markers, and ultimately identified these cells as a subset of cDC2 cells. We conducted experiments showing that the FR phenotype appears to be a transient state that APCs pass through on a 1 hr time-scale rather than a distinct cell subset. We tested whether these cells alone were sufficient to mediate the response of the remaining APCs in a culture – concluding that they can trigger neighboring APCs via paracrine signaling. We examined this further and found that for an adoptive transfer experiment of DCs, isolating only the 5% of FR cells and transferring them resulted in nearly the same level of adaptive immunity as transfer of the whole culture. This study shows how the transient state of FR are important players in innate immune activation and have potential for therapeutic targeting.

## Results

### TLR Agonist Conjugated MP For Isolation of FRs

The primary hypothesis of this paper is that there exists a rare and TLR responsive DC state, which we refer to herein as first responders (FRs). These cells assist and amplify innate activation and adaptive responses via paracrine signaling and direct action. The first responder state is characterized by (1) rapid accumulation of micron sized TLR agonist coated particles far exceeding statistical probability, (2) rapid, high-level expression of inflammatory cytokines which activate bystander cells, (3) a preliminarily identified set of cell surface markers which identify the transient cell state in naïve populations. From initial experiments, evidence supports that most cells sample this state rather than it being a unique phenotype and that approximately 5% of the population is in a first responder state at any given time. First responders represent a subset of the DC response which we have identified can be targeted selectively for adaptive response to a pathogen (Figure 1A). We conjecture that perhaps the FR state has not been characterized previously due to the low percentage of cells, transience and accompanying difficulty in identifying distinguishing cell-surface markers. Our first objective of this study was to develop a method to reliably isolate FRs from a heterogenous population of DCs. As FRs respond to TLR signaling^2^, we utilized a TLR conjugated micro-particles (MP) system to isolate FRs from other DCs. In previous work, we had developed a system for studying how particles interact with innate immune cells. ^3^ We synthesized 2 µm diameter FITC labeled polystyrene MPs and then coated them with a silicon-siloxane coating to reduce non-specific immune reactivity (Figure S-1). The MPs were surface conjugated with one of 5 different TLR agonists (for TLR2, 4, 5, 7 and 9) using maleimide-thiol chemistry (Table S-1, Figure S-1). Using previous established methods, we calculated an approximation of the number of TLR agonists per MP using a BCA (bicinchoninic acid assay, quantifying the number of amide bonds) for the three agonists with amide bonds (LTA, LPS and FLA) and calculated approximately 1-10 million TLR agonists per MP (Figure S-1E). We then incubated the MPs with innate immune cells and used particle uptake to isolate FRs from non-FRs (nFRs) (Figure 1A).^12^

**Figure 1.**
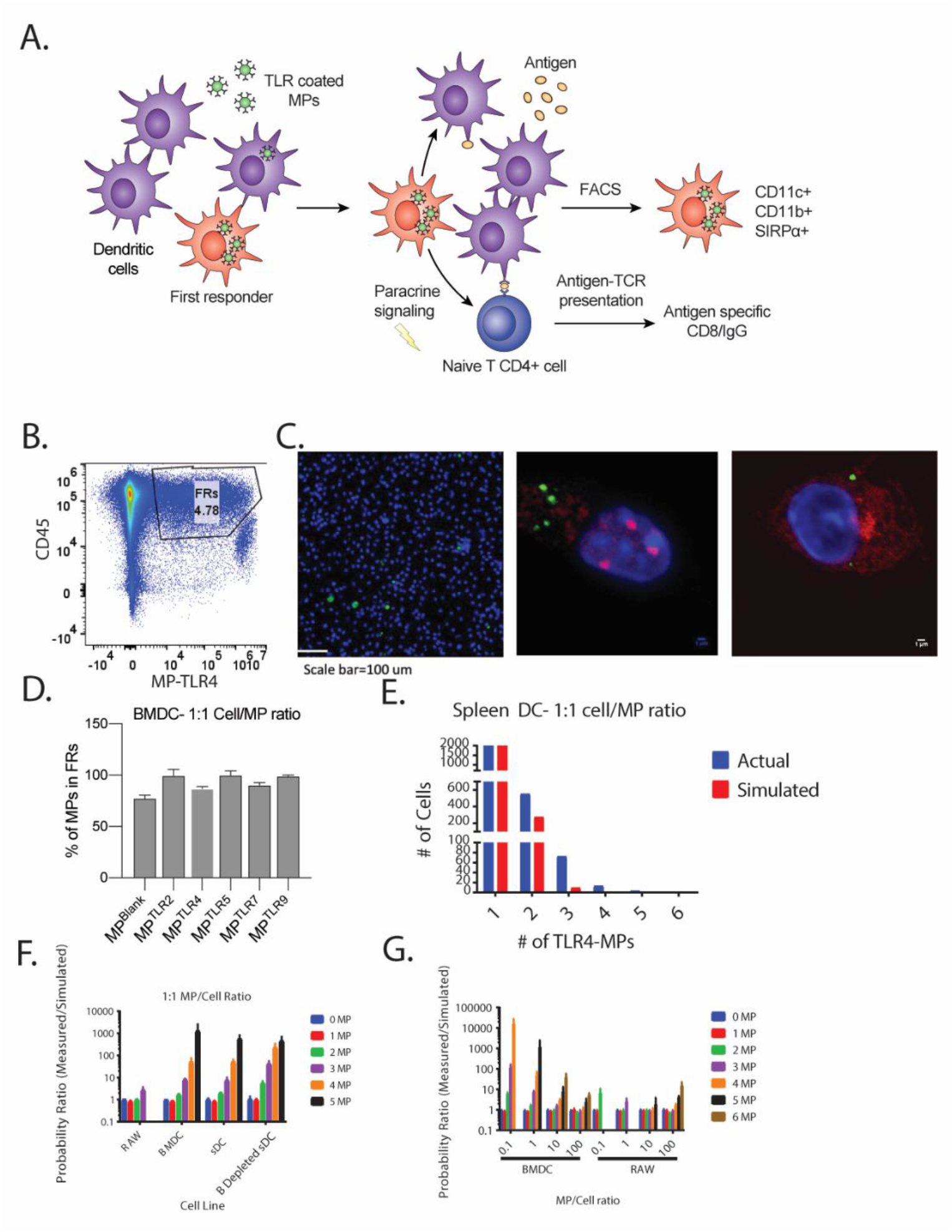
Identification of First Responder Cell State With Statistically Improbable Uptake of TLR coated MPs. (A) Schematic figure illustrating the overarching hypothesis of the study. FRs uptake more TLR coated MPs, activating neighboring APCs via paracrine signaling. (B) Flow cytometry of mouse spleenocytes incubated with MP^TLR4^ 1:1 for 30 minutes. FRs are defined as top 5% of MP signal. (C) Confocal microscopy images of BMDCs incubated with MP^TLR4^ 1:1 for 30 minutes. Blue channel (DAPI), green channel (FITC MP^TLR4^), Red channel (NF-kB). Left image: wide field view of BMDCs and MPs (no NFkB staining). Middle Image: FR with NFkB staining, Right-nFR with NFkB staining. (D) Percentage of MPs in FR cells. 100K BMDCs were incubated with 100K of varying MP formulations for 15 minutes, washed and then analyzed via ImageStream. The number of MPs uptaken by each cell was determined using the “spot counter” function on the IDEAS software and the percentage of all MPs in FRs was calculated. (E) Spleen derived CD11c+ cells (sDCs) were incubated with MPs at a ratio of 1:1 for 15 mins, stained for CD11c+ and analyzed via Imagestream similar to part D. The number of cells was normalized to 10k total MPs uptaken per sample. The number of cells with 1-6 MPs in these samples (actual) were compared to a random Poisson distribution (simulated). (F) This analysis was repeated for RAW cells, BMDCs, and B cell depleted CD11c+ spleenocytes and a ratio of the actual/simulated was determined. (G) Repeated analysis from part F but varying the ratio of MPs to cells. All experiments were performed in biological triplicates, error bars indicate ±SD.

### FRs Uptake Significantly Greater Number of MP

We initial identified the first responder phenomenon by happenstance. In previous work, we were calculating the number of agonists on a particle which activate a cell.^13^ During the course of these experiments, we noticed a very large statistical anomaly - a small percentage of DCs had a disproportionally high MP signal that deviated greatly from a normal (Poisson) distribution. We expected any immune cell would phagocytose MPs and especially MPs conjugated with TLR agonists, but based on a random distribution, it is expected to be a normal distribution of cells per MP assuming all DCs are homogeneous in their MP uptake properties.^14^ However, when we incubated a heterogeneous cell line of either bone marrow derived dendritic cells (BMDCs) or spleen derived DCs (sDC) or a homogeneous control macrophage cell line (RAW) with MPs, we observed a skew in higher fluorescence intensity for the heterogeneous cell lines but not for RAW cells (Figure 1B, Figure S-2, Figure S-3 for characterization of BMDCs). This effect was also seen via confocal microscopy, where MPs aggregate mainly in a select few BMDCs. At the same time, most of the TNF-α secretion was corresponding localized to the same few cells (Figure 1C). This result was supported by observations published previously by Miller-Jensen and co-workers.^2^ To further quantify this shift, we used Image Stream flow cytometry (ISX) to count the number of particles phagocytosed per cell. We observed that >90% of all MPs (an LPS conjugated MP) were phagocytosed by <5% of all BMDCs (Figure 1D). This cutoff of top 5% of FITC signal was used to define FRs in all following experiments in this study.

To explore this phenomenon further, we analyzed sDC or BMDCs via ISX and observed an unusually high number of cells with >2 MPs per cell. When compared to a standard “random” Poisson distribution, there was a larger number of cells with >2 MP for sDCs and BMDCs but not for the RAW control cell line (Figure 1E, Figure S-4). We were intrigued by this dramatic shift for a small number of MPs in the heterogeneous cell lines and hypothesized if this increase could be explained by normal statistical variance or if there was something distinctive about this sub-population of cells. To do this, we further compared the standard Poisson distribution to our measured result, normalizing both the number of cells analyzed and the total number of MPs uptaken for each experimental condition and calculated a ratio of measured probability a cell has uptaken a certain number of MPs over the “simulated” probability based on the Poission distribution (Figure 1F). For heterogeneous cell mixtures (BMDCs, sDCs or sDCs depleted of B cells), there is an increase in the probability ratio for cells with >2 MPs, but not for RAW cells (Figure 1F). This result was also observed in all other MP formulations (Figure S-5).This result is unexpected, as similar studies using TLR coated MPs and BMDCs did not observe high concentration of particles in certain cells.^15^ We believe this to be an effect of MP to cell ratio during incubation, as previous studies used a MP to cell ratio (>10) whereas our experiments used a lower 1:1 MP/cell ratio. We compared four different MP to cell ratios in a similar experiment to part F and determined that this increase in MP uptake for FRs is only seen in lower MP/cell ratios and at higher ratios (10, 100 to 1 MP to cell) had >15% of MPs contained in FRs (Figure 1G, Figure S-6). It is likely that at higher ratios, MPs will “saturate” all cells and obscure the increased uptake of MPs in FRs, which might explain why the FR phenomena was not observed previously.

### FRs are primarily Central Dendritic Cells Type 2 (cDC2)

We sought to phenotype FRs to determine if they are an already described phenotype of APCs. We phenotyped spleenocytes incubated with MP^TLR4^ to determine if FRs match one of the major DC subsets, with a focus on cDC1 or cDC2 (see figure S-7 for gating strategy). When isolating FRs, we observed an increase in the number of cells in the cDC2 compartment (SIRPα+, CD11b+) for FRs compared to non-FRs (Figure 2A). When gated on the DC compartment (CD11c+, MHCII hi), over 70% of FRs were also cDC2, while only 38% of DCs are naturally cDC2. The increase in cDC2 percentages were also seen when gated on all CD45+ cells and for BMDCs (Figure2A). We observed a similar increase in cDC2 incidence for FR populations with spleenocytes incubated with other TLR agonist conjugated MPs, indicating that the FR state is primarily a part of the cDC2 populations (Figure 2B) This same result in seen in resident skin DCs, where mouse footpads injected with MPs primarily are uptaken by resident cDC2 cells (Figure S-8).

**Figure 2.**
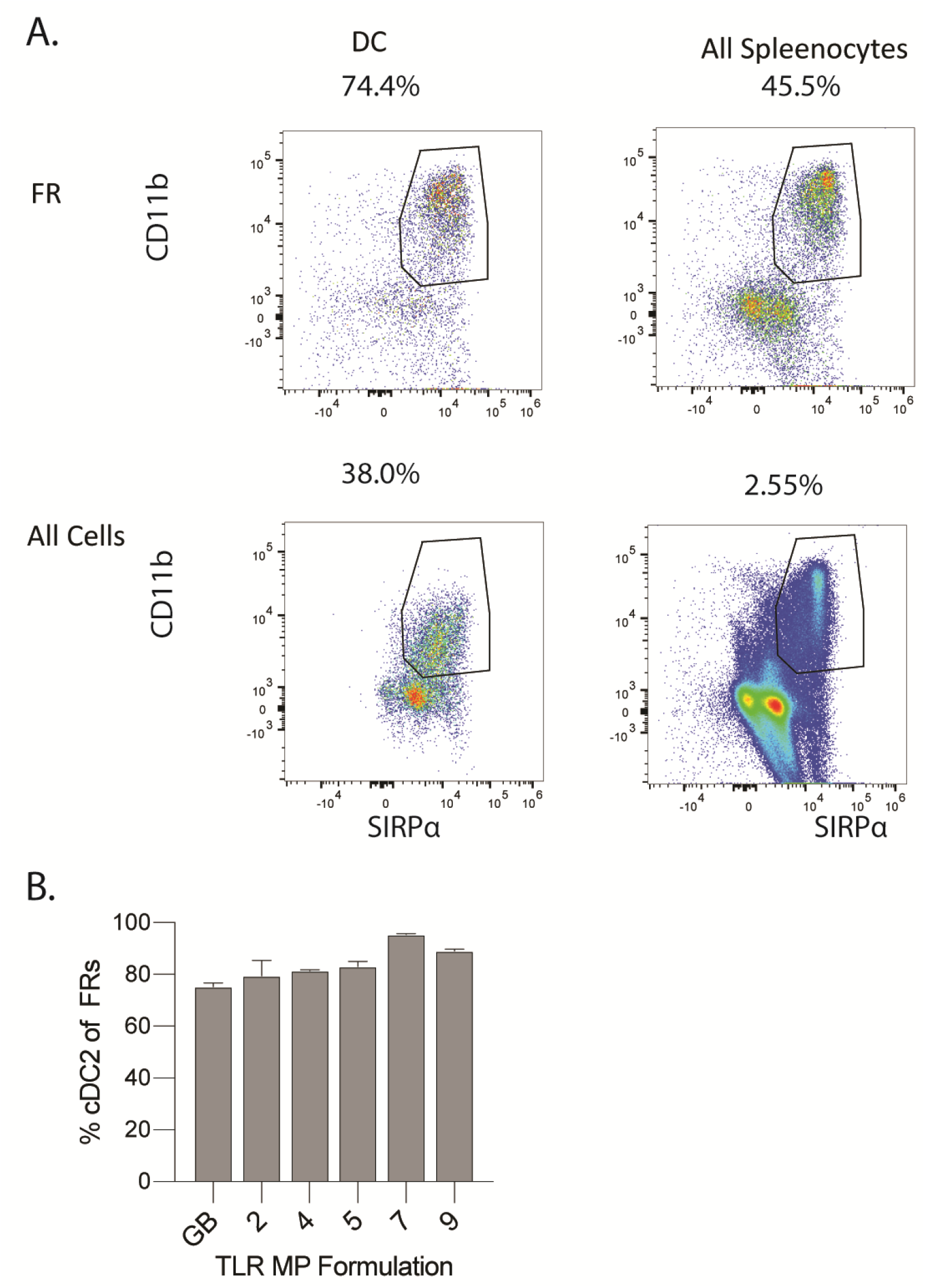
FRs are primarily central dendritic cell 2 (cDC2) class. (A) Mouse Spleenocytes were isolated, incubated with MP^TLR4^ and phenotyped via Aurora flow cytometry. Representative flow plots for DC (CD11c+, MHCII hi) and/or FR (top 5% of MP signal). (B) Graph showing percentage of FRs that were also cDC2 given various MP stimulation similar to part A. All experiments were performed in biological triplicates, error bars indicate ±SD.

### FRs have increased TLR activation, coordinate local APC activation via paracrine signaling and are temporally controlled

After phenotyping the FRs and determining that they are primarily part of the cDC2 population, we were interested in studying their cytokine profiles and kinetics of activation. Since the FRs are the first cells to phagocytose MPs with PRRs, we hypothesized that the FRs could also play a large role in initiating the innate immune response. As such, we wanted to gain insight into the nature of their immune response. There were two major questions that we wanted to answer. 1) How does the immune response of FRs differ from that of all the other cells which we refer to as non-first responders (nFRs)? 2) What is the kinetics of the FR immune response and how do these kinetics influence bulk immune activation *in vitro*? We sought to validate these questions in vitro and then, if either yielded results, to explore them in vivo. To probe the first question, we investigated the cytokine expression of the FRs compared to the nFRs. Because we had observed that FRs produce high levels of cytokines, we limited intercellular cytokine signaling, by preincubated naïve BMDCs with Brefeldin A. Then we stimulated 100k BMDCs with MP^TLR-X^ (x = blank, 2, 4) in a 1:1 ratio for 1 hr, washed and incubated with Brefeldin A for 16 hrs and analyzed the cells with Imagestream flow cytometry (ISX) to determine TNFα expression. TNFα is a strong proinflammatory cytokine, and the FRs had a two-fold increase in TNFα relative to nFRs indicating that FRs produce higher levels of proinflammatory cytokines than the nFRs.^16^ (Figure 3A). We also observed this increase via confocal microscopy (Figure S-9). To further probe the cytokine profile of the FRs, we looked at TNFα release in FRs and nFRs populations after stimulating naïve BMDCs with MP^TLR-4^ at 1:1 ratio. The FRs and nFRs were isolated via FACS, washed, and resuspended in media at a concentration of 1 million cells per mL. After 1 h, the supernatant was collected and the TNFα was measured via cytokine bead array (BD Biociences). The supernatant was collected after 1 h, and similar to Xue *et al*., we observe that a small percent of cells, in our case the FRs, are responsible for significantly more TNFα secretion than the nFRs.^2^ in the nFR cell population we observe that the TNFα secretion has relatively small increase (∼ 50%) relative to baseline. However, the FR population secretes significantly more TNFα than the nFR population. The measured TNFα levels secreted by the FRs is 1406 pg/mL, almost a 10 fold increase from the untreated population (Figure 3B). This result indicates that the FRs, on average, secretes over 6 times the amount of TNFα per cell compared to the nFRs. From these data, we hypothesized that the FRs could be key for stimulating immune activation in neighboring cells. We also noted that IL-1β was not increased in FR populations, indicating that our MPs do not activate inflammasome activity (Figure S-10).^15^ We further confirmed this lack of inflammasome activity by demonstrating that MPs do not escape lysosomes in FRs (data not shown).

**Figure 3.**
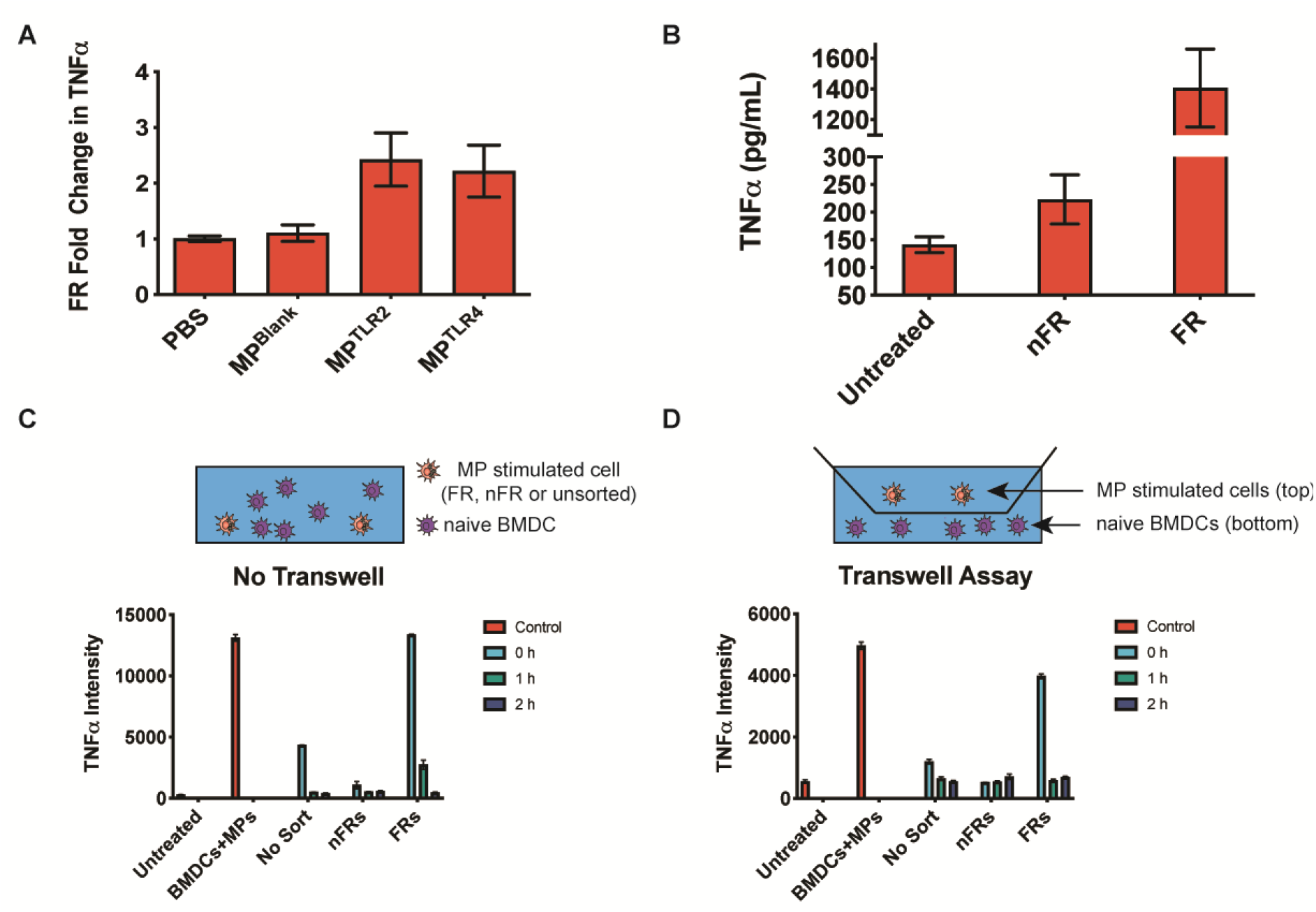
FRs are sufficient and necessary for activating populations of naïve BMDCs *in vitro*. A) BMDCs incubated with Brefeldin A, (1 ug/mL) MP^TLR-X^ (x = blank, 2, 4) were added to 100k BMDCs. After 6 h, the TNFα levels were analyzed via ISX. FRs express higher levels of TNFα than nFRs. B) BMDCs were stimulated at a 1:! ratio with MP^TLR-4^ for 15 min. The FRs and nFRs were isolated, washed, and resuspended at 1 million cells/mL in 10% HIFBS in RPMI for 1 h. The supernatant was collected and profiled via cytokine bead array (BD Biociences). FRs secreted 1406 pg/mL of TNFα – 6.4 times more TNFα per cell when compared to nFRs –. C) Naïve BMDCs were stimulated at 1:1 ratio with MP^TLR4^ and isolated the FRs and nFRs via FACS. After 0, 1, or 2 h incubation, FRs, nFRs, and unsorted BMDCs were added to 1 million naïve BMDCs in a 1:10 ratio. After 16 h, TNFα intensity was measured, and there was high TNFα expression in the bulk population when the FRs were added at 0 h incubation time. There was a background levels of TNFα expression when nFRs were added, regardless of timepoint. D) A similar experiment to C, but the naïve BMDCs were plated on the bottom section of a transwell assay and the MP-stimulated FRs, nFRs, and unsorted BMDCs were plated on top of the membrane. Similarly, naïve cells subjected to FRs express significantly higher levels of TNFα at 0 h incubation. All experiments were performed in biological triplicates, error bars indicate ±SD.

After observing the high TNFα levels produced by FRs, we sought to determine if FRs can induce a pro-inflammatory response in an *in vitro* population of naïve cells and also the timeline of when FRs signal to neighboring cells. To test these questions, we activated naïve BMDCs with MP^TLR4^ and isolated the FRs and nFRs via FACS. We isolated cells immediately after sorting – designated as t = 0, 1, or 2 h after incubation. FRs (top 5% of MP signal), nFRs (bottom 90% of MP signal), or unsorted BMDCs were washed after sorting and then added back to a fresh-culture of 1 million naïve BMDCs from the same pool of cells in a 1:10 ratio. After 16 h, immune activation was determined by measuring the intensity of TNFα and CD40, a costimulatory molecule and marker of immune activation.^17^ We only observed high activation in BMDCs that were subjected to FRs at 0 h. From this, we conclude that the FRs are necessary for bulk *in vitro* activation of BMDCs. Interestingly, we measured only background expression of CD40 and TNFα when the FRs were incubated for 1 or 2 h before addition to the naïve BMDCs, indicating that the FR response very short-lived. Furthermore, in the cultures stimulated with nFRs, only background levels of TNFα and CD40 were measured, indicating that nFRs are not sufficient to stimulate *in vitro* immune response (Fig 3C-3D, Figure S-11).

After determining that FRs are a critical component of obtaining an in vitro response, we wanted to investigate whether the activation relied only on soluble factors. To test this, we repeated this experiment using a transwell membrane where the FRs, nFRs, and unsorted BMDCs were plated on top of the membrane and the naïve cells were plated below the membrane. Similarly, we observed higher levels of CD40 and TNFα at 0 h, only when FRs were added to naïve cells (Fig 3D, Figure S-11). These data indicate that secreted proteins working via paracrine signaling, driven by FRs, play an important role in the bulk activation of BMDCs, but only on a short, < 1 h timescale after initial FR stimulation. It is important to note that this activation only occurs when the FRs are immediately added to the naïve BMDCs. If the FRs sit for > 1 or 2 h, the responses of the whole *in vitro* culture give responses similar to unstimulated cultures. Most intriguing, we observed that the FR state is periodic and renewable. When we isolated nFRs, upon further culture, these isolated cells would then product FR-like cells which could be used isolated, stimulated and restimulate a naïve culture of BMDCs in an identical manner to initial FRs. However, this phenomenon only occurs after nFR rest for several hours after isolating the first set of FR (Figure S-12) indicating that the FR state renews on a multi-hour period. From these experiments, we conclude that FRs are both sufficient and necessary to initiate immune activation *in vitro* in BMDCs but can only perform this bulk stimulation on a short timescale after their initial activation.

### FRs are necessary and sufficient for generating antigen specific responses via BMDC adoptive transfer

After determining how FRs coordinate innate immune activation and their kinetics, we wanted to observe the impact of FRs on adaptive immune responses. To understand this further *in vivo*, we performed an adoptive transfer of MP treated BMDCs with C57BL/6 mice in combination with the antigen ovalbumin (OVA). We had observed in vitro that BMDC FRs have increased expression of MHCI and II and present more OVA, so we hypothesized that they might also play a larger role in antigen presentation and formation of adaptive immune responses (Figure S-13) To make a preliminary assessment, we compared if BMDC FRs could recapitulate part or all of the stimulation of an adaptive response of a larger collection of BMDCs in an adoptive transfer experiment. We hypothesized that if the FRs were controlling a larger portion of the response, then removing the other cells would have a minimal impact on the final adaptive response. BMDCs were incubated with MP^TLR4^ on a 1:1 cell/MP basis for 30 minutes, washed and then concentrated at 30 million cells/mL (or 3 million in the case of FRs) with 10 µg/mL of OVA. To accomplish this goal, FR isolated and washed from a larger population of BMDCs were injected into both mouse footpads at 1 million cells per footpad for unsorted group or nFRs and at 100k per footpad for FR groups (Figure 4A). All experimental groups were treated with OVA except for PBS controls. We previously determined that BMDCs injected into the footpad migrate to the popliteal lymph nodes and confirmed that these cells initiate immune responses (Figure S-14). After 14 days, adoptively transferred mice were sacrificed and their sera was sampled for OVA specific IgGs.

**Figure 4.**
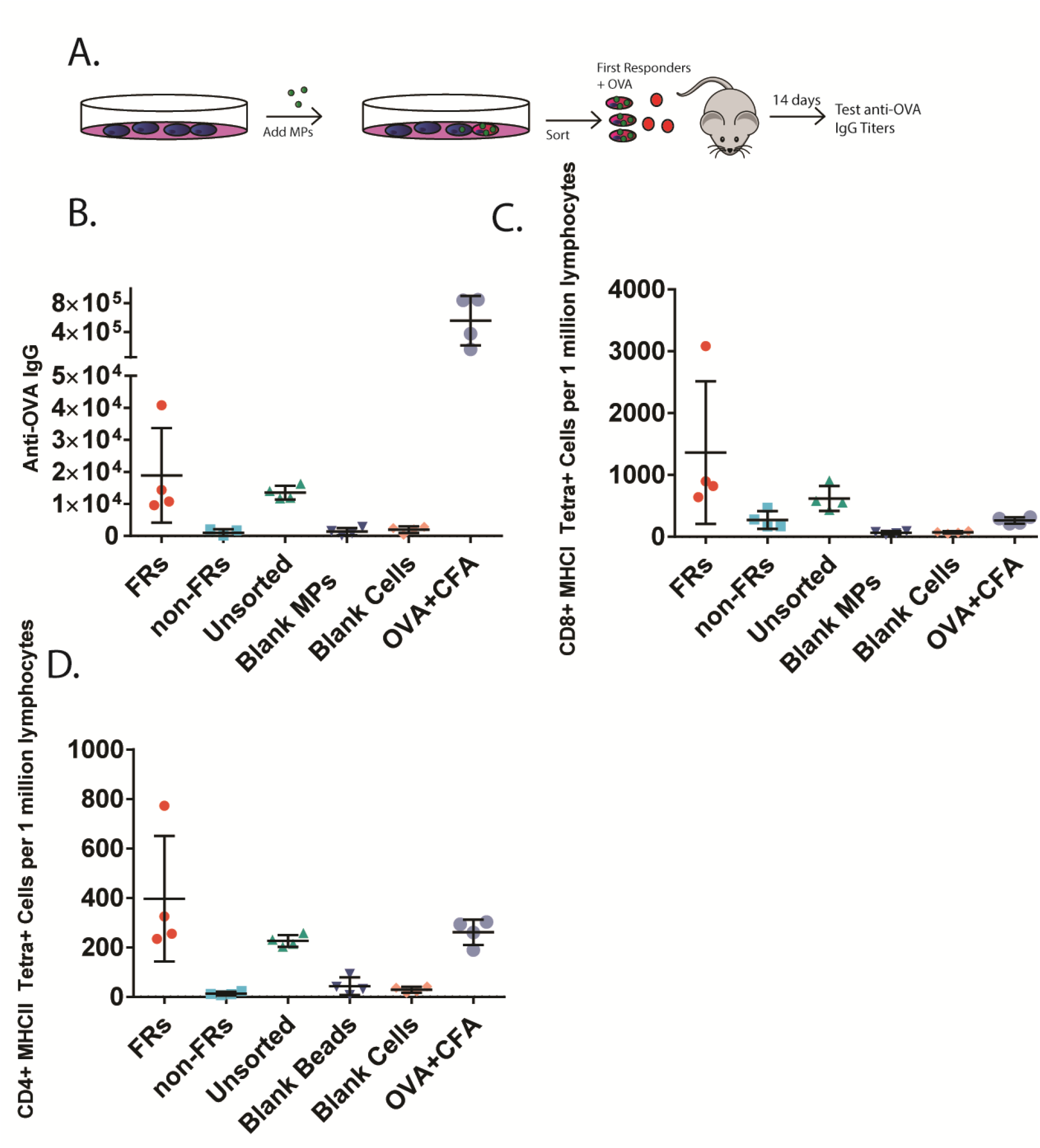
FRs are necessary and sufficient for triggering adaptive immune responses in vivo. (A) Schematic of FR adoptive transfer experiment. BMDCs were incubated 1:1 with MP^TLR4^ for 30 minutes, washed and sorted into FR or nFR populations. Both hind leg footpads of a C57BL/6 mouse were injected with either 100K FRs, 1 million nFRs, 1 million unsorted MP incubated BMDCs, untreated BMDCs, CFA (positive control) or PBS (negative control). All samples except PBS contained OVA (10 ug per footpad). (B) anti OVA IgG titers 14 days post injection. (C) CD8+ T cells positive for a MHCI tetramer to the major MHCI epitope OVA 257-264. (D) CD4+ T cells positive for a MHCII tetramer to the major MHCII epitope OVA 323-339. Each group had N=5 mice, error bars indicate ±SEM.

### Popliteal lymph nodes were disaggregated and cells analyzed

Mice injected with isolated FRs had a >10-time increase in anti-OVA IgG titers over mice injected with BMDCs not treated with MPs (Blank Cells) (Figure 4B, p<0.05). This increase is particularly striking because FRs were injected with 10-fold fewer cells than the blank cell group and blank MP control group. Furthermore, when 1 million non-FRs were transferred in place of the 100k FRs, very little OVA-specific IgG was measured. Adding to this, when the unsorted mixture of 1 million FR and nFR was transferred a response nearly equivalent to the FRs was measured, indicating that FRs are necessary to trigger adaptive immune responses in an adoptive transfer. To ensure that particulates did not influence antibody responses, identical controls were performed with MPs containing no TLR agonists prior to transfer resulting in no IgG production. Similar trends between treatment groups were also seen in the production of OVA specific CD4 T cells for both the major MHCI epitope (OVA 257-264) and the MHCII epitope (OVA 323-339) and for CD8 T cell responses (Figure 4C, 4D, Figure S-15).^18^ It is important to note that while FRs seem to be critical for BMDC mediated activation via adoptive transfer, the restriction of the APCs used to activate adaptive immunity may have reduced overall immune response. In contrast, a positive control using mice injected with OVA and Complete Freud’s adjuvant (CFA) which stimulated only native APCs resulted in 10-fold higher IgG titers. From this data, we conclude that in adoptive transfer experiments FR contribute an out-sized proportion of the response in antigen presentation and DC mediated induction of adaptive immunity given their very small population numbers pointing to the possibility of a large role for in vivo responses.

### FRs have a unique temporal transcription profile

Given the importance of FRs for developing adaptive immune responses, we sought to further characterize FRs activity via whole transcriptome sequencing. We incubated BMDCs 1:1 with MP^TLR4^ for 15 minutes, washed, isolated FRs (the top 5% of MP signal) from the nFRs (bottom 90%) using FACS, and incubated for either 0 (immediately after sorting), 0.5, 1, 2 and 4 hrs and then isolated mRNA using a commercially available kit (Illumina) and sequenced the mRNA with the aid of the UChicago’s genomics core. The mRNA sequences were aligned to the mouse transcriptome and two-fold mRNA upregulation compared to an untreated BMDC control sample. We noticed that at the 0 h timepoint, transcription levels for several inflammatory cytokines, such as TNFα, INFβ, CXCL1 and IL1β and many others (Figure 5E) were highly upregulated in FRs but not upregulated in nFRs (Figure 5A-E). In later timepoints, these cytokines mRNA levels decreased for FRs but increased for nFRs. For example, TNFα mRNA levels dropped to below baseline after 1 h for FRs but peaked for nFRs at 1 h (Figure 5A). This further confirms our hypothesis that FRs have a burst release of inflammatory paracrine signaling cytokines immediately following TLR engagement, which simulates neighboring cells at later timepoints. This observation agrees with our previous observation that FRs accumulate the majority of TLR-bearing particles thereby providing the only possible point for TLR-mediated activation of innate immunity.

**Figure 5.**
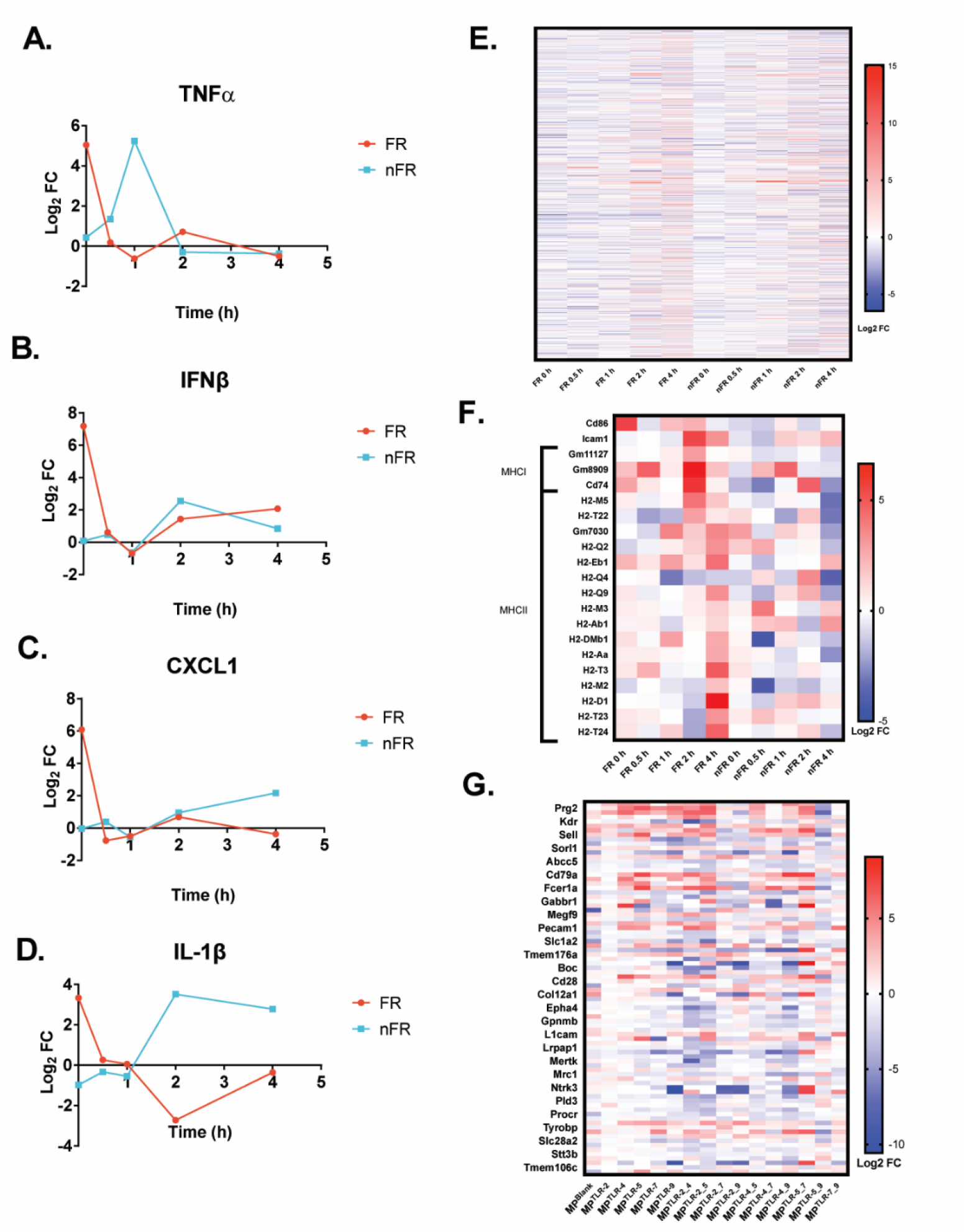
FR mRNA Analysis. BMDCs were incubated with MP^TLR4^ 1:1 for 15 mins, washed and immediately sorted into FRs (top 5% of MP signal) and nFRs (bottom 90% of MP signal). The mRNA from these cells were isolated immediately off sorter (0 hr) or incubated at 37°C for 0.5, 1, 2, or 4 hrs and isolated. cDNA was generated from the poly A mRNA using a commercially avaliable kit from Illumina and sequenced. Genes were aligned and two-fold upregulation calculated by comparing to a non-treated BMDC control. Timecourse fold change of the following cytokines were plotted, (A) TNFα, (B) INFβ, (C) CXCL1 and (D) IL-1β. (E) Heatmap of log2 fold change of mRNA for genes with known immune function over timepoints. F) Heatmap of log2 fold change in the mRNA for genes with known antigen presentation function. Genes with pval <.05 and 2 fold differential expression for at least one of the timepoints were included. (G) BMDCs were incubated with varying MP formulations for 16 hrs with Brefeldin A treatment, washed and sorted into FRs and nFRs as in part A-E and mRNA immediately isolated, transformed into cDNA, sequenced and genes aligned. A list of upregulated mRNA genes in FRs that correspond to surface receptors is shown in a heatmap. All experiments were performed in biological triplicates with significant upregulation determined only if the pval < <0.05.

It is surprising that this TLR-mediated immunity is then amplified so widely by the nFRs in such a short time. Also, of interest is that the timing of transcription response for the nFR more closely match the responses normally associated with a response to TLR stimulation for a population of BMDCs.^19^

In addition to the cytokine expression, we wanted to determine whether the primary function of the FRs immune response is limited to the quick cytokine burst, or if it extended to other aspects of a typical immune response, such as antigen presentation. A list of antigen presenting genes were acquired through the Mouse Genome Informatics database, and genes with 2-fold change in expression and pval < .05 for at least one time point were analyzed. Overall, we observe more upregulation for MHCI and MHCII genes in the FRs than we do in the nFRs (Figure 5F). FRs upregulated antigen presenting genes at 2 and 4 hr while nFRs did not demonstrate upregulation. Likewise, the MHC co-stimulatory molecule CD86 is upregulated in 0 hr time point in FRs whereas no upregulation is seen in nFRs. Furthermore, as highlighted earlier, we have seen a 2-fold increase in MHCI and MHCII protein expression in FRs 15 mins after sorting (Figure S-13). From this data, we expect the FRs to not only be responsible for bulk immune stimulation, but also to be important in the initial antigen presentation as well.

In order to find unique identifying proteins of the FR state, we analyzed the transcriptional response for upregulated transcriptions of FR cells corresponding to potential surface proteins. One unique challenge for identifying surface protein upregulation in such a time-sensitive and dynamic cellular state as FRs is determining the appropriate timepoint for observing mRNA expression.^20^ In light of this, we preliminarily incubated BMDCs with varying MPs for 16 hrs in Brefeldin A then sorted FR and nFRs and isolated mRNA. This timepoint was chosen for two reasons: (1) brefeldin A has been shown to cause mRNA accumulation over time and (2) while it is unlikely that mRNA for already expressed proteins would be upregulated at the time of sorting, we hoped to observe upregulation on the next FR “cycle”.^21^ Additionally, we did not want the unique identifying proteins to depend on the TLR agonist. Thus, we incubated BMDCs with MP^TLR-X^, where x is various single or dual TLR agonist(s) attached to the MP for 16 h and isolated the FRs and nFRs for sequencing via FACS. To find unique proteins, we looked for the genes that were most consistently upregulated amongst these dosing conditions (2-fold increase in expression and pval < .05) when comparing the transcriptional profiles of the FRs to the nFRs. The data showed upregulation of several surface proteins of interest and while further study would be necessary to identify an optimal timepoint, this provided a list of potential targets of over 25 potential proteins for FR identifying markers (Figure 5G).

### FRs express unique and identifiable surface proteins

One initial limitation in our identification of FRs was the lack of identifying cell-surface markers – allowing them to be isolated from other DC cells. This problem is compounded by their transience and low-percentage within the cDC2 and DC population. However, our mRNA analysis pointed out several proteins which suggested distinctive cell-surface markers. To identify candidate protein for identifying FRs via a flow cytometry panel, we selected proteins: (1) whose mRNA was upregulated in FRs compared to nFRs using all or most of the MP formulations, (2) were surface expressed and had commercially available high affinity antibodies, (3) had published low expression in other cells types and (4) were well characterized in the literature. We believed that these 4 characteristics would not only narrow our search, but also provide viable candidates to readily identify the FR state. Allowing target proteins that were already well characterized also opens the potential to enable targeting FRs for therapeutic purposes. We tested potential candidates using flow cytometry, incubating with MPs for 15 mins to isolate FR using our established method then washing and staining for candidate markers to see test if the candidate markers identified the same cells. Using BMDCs and our mRNA analysis, we narrowed the many candidates that were highly upregulated to just a few by applying a second set of conditions including eliminating non-surface markers or markers that are present in many types of immune cells such as CD20 or FTL-3 (Figure S-16). Given the presence of some mRNA corresponding to what are canonically considered B cell markers, we also validated that FRs do not express actually common B cell markers such as CD19 and B220 (Figure S-17). From this selection, we identified three markers of interest, DAP12, PRG2 and TMEM176A, that were upregulated 3-7 time between FRs and nFRs in BMDCs (Figure 6A). Furthermore, they were even more highly upregulated in mouse spleenocyte CD11c+ cells and in B cell depleted spleenocyte CD11c+ cells (Figure 6B, C). Importantly, there was not a high fold change in control samples using a homogeneous cell line (RAW), confirming that the observed increase of expression is not an artifact of our analysis but rather due to the heterogeneity of the cell population themselves (Figure 6D).

**Figure 6.**
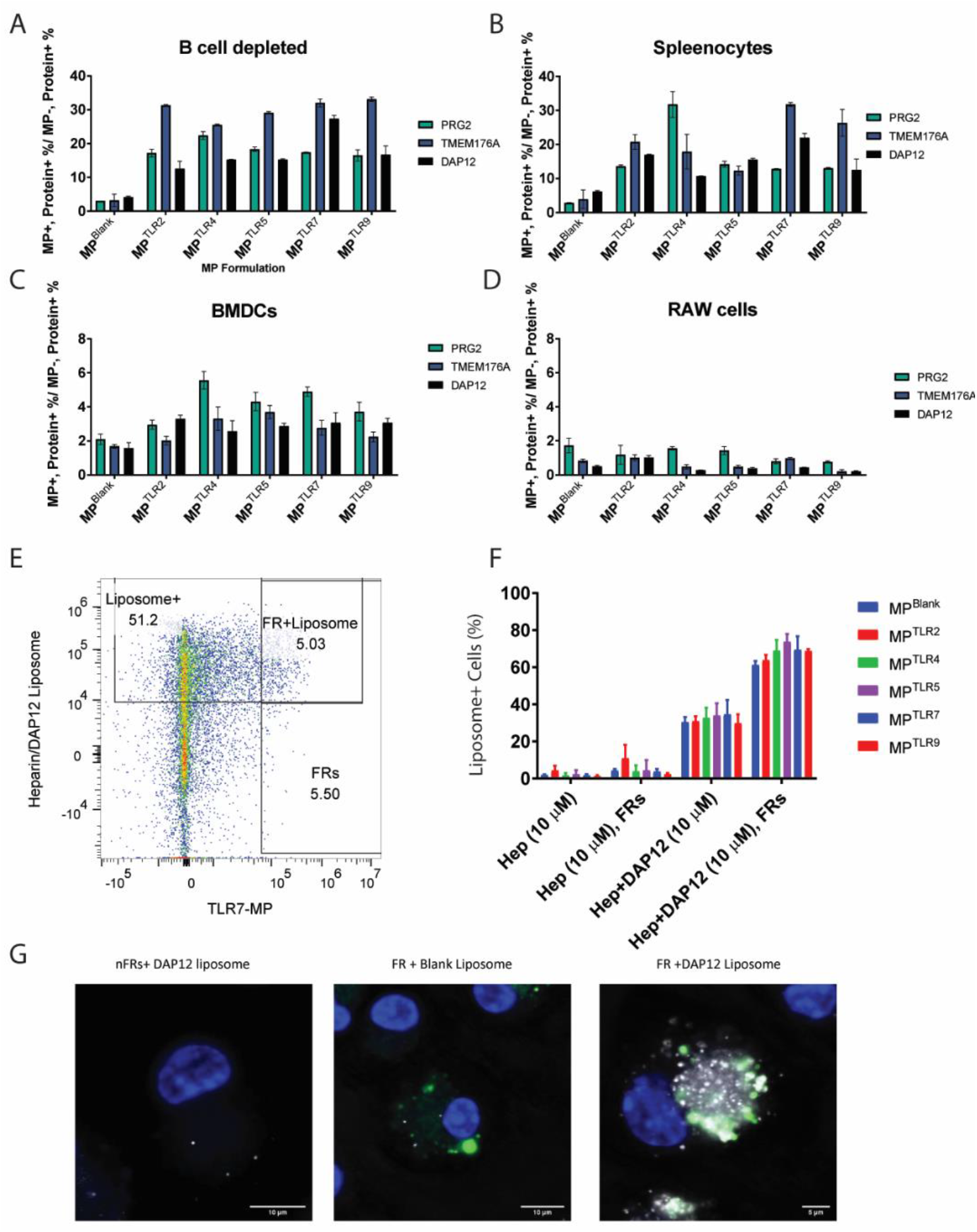
Identification of Cell-Surface Markers of FR and Targeting of FRs with Liposomes. (A) 1 million BMDCs were incubated with 1 million various MP formulations for 15 minutes, then stained for CD11c and either PRG2, DAP12 or TMEM176A and analyzed via flow cytometry. % Positive Ratio represents the ratio of PRG2, DAP12 or TMEM176A positive cells in the FRs divided by % positive cells in the nFR population of the CD11c+ cells. (B) Similar to part A but using mouse spleenocytes or (C) B cell depleted mouse spleenocytes or (D) RAW cells. (E) 1% DAP12 peptide, 10% heparin-lipid loaded, DiD labeled (0.01%) 200 nm diameter liposomes were incubated at 10 uM total lipid concentration in combination with 1 million MP^TLR4^ with 1 million spleenocytes for 30 mins. Cells were washed and stained for CD11c and analyzed via flow cytometry. Plot is a representative example of gating strategy for CD11c+ cells to define Liposome+ and FRs. (F) Graph demonstrating the percentage of cell population that were liposome+ for various MP formulations. (G) Representative confocal images of BMDCs incubated with 10 uM total Lipid of targeted liposome and MP^TLR7^. Left-nFR (with no MP signal) with targeted liposome, Center-FRs incubated with 10 um of non-targed liposome, Right-FRs+ 10 uM of targeted Liposome. All experiments were performed in biological triplicates, error bars indicate ±SD.

Cells in the FR state were also more heterogeneous and dynamic that we first anticipated. There appears to be increasing levels of upregulation of DAP12, PRG2 and/or TMEM176A depending on the TLR agonists conjugation to the MP. Unsurprisingly MP^Blank^ had the lowest fold increase, but there were very high increases for PRG2 with MP^TLR4^ and DAP12/TMEM176A with MP^TLR7^. This result suggests there may be different subpopulations of the FR state which correlate with different TLRs or TLR agonists, although further research would be required to validate this hypothesis. Nevertheless, these three markers (1) repeatedly showed upregulation for each TLR-bearing MP, (2) are surface expressed, (3) are present during immune activation and antigen processing, and (4) are not highly expressed in other cells. From these four points, we considered them viable markers which warranted further study.^22–24^

### FRs can be targeted by their surface proteins

After confirming that the FRs both had unique uptake, markers, and contributed to a large degree of an adoptively transferred BMDC stimulation, we sought to examine how much of an in vivo response was mediated by FR state. To accomplish this, we reasoned, we would need a method to target and ablate the FR state during the short period when each cell was transitioning through it. To target the FRs, we used a multivalent approach, combining ligands for several of the markers we had identified through flow cytometry. To bind DAP12, we searched the literature and found a selectively binding nonamer peptide, GFLSKSLVF. To bind PRG2, we identified heparin which has moderate affinity.^25,26^ In order to test if these targeting elements would identify FRs, we used a well characterized liposomal system to insert both a DAP12 and Heparin conjugated lipid into 200 nm DSPC liposomes. We synthesized and purified lipid conjugated versions of the DAP12 and heparin polymer(Figure S-18).^27,28^ We then generated liposomes using the membrane extrusion technique (200 nm filters, see methods for additional details on liposome synthesis and composition). By inserting the targeting elements during formulation, we generated several ratios and compositions of the two targeting elements. We determined that the optimal ratio of targeting elements in the liposome was 10% heparin-lipid and 1% DAP12 lipid (Figure S-18). We then coincubated a fluorescently labeled version of the heparin/DAP12 liposome with BMDCs or spleenocytes for 15 mins, and washed to remove excess liposomes. After staining cells with potential FR-identify liposomes, we incubated cells with TLR conjugated MPs for 30 minutes, washed and observed uptake via flow cytometry for MPs and liposomes. (Figure 6E, F). Only when the liposomes contained both heparin and DAP12, did we observe a significantly higher percentage of liposome positive cells for the FR population, indicating that targeting both receptors was necessary (Figure S-19). This was also observed via confocal microscopy, where BMDCs that had high MP signal also had high liposome signal only in the targeted formulations but not for BMDCs with no MP uptake or BMDCs incubated with blank (non-targeted) liposomes (Figure 6G). This data indicates that we can reliably target FR cells with the heparin/DAP12 peptide liposome formulation and that they match the population we previously identified via our microparticle experiments. With the surprising ability of these liposomes to identify FRs, which we conjecture is helped by their active sampling of their environment, we were intrigued if this targeting could be used to enhance FRs ability to amplify immune responses.

### In Vivo Validation of FR Ablation and Targeting

After demonstrating that FRs can be reliably targeted via DAP12/Hep loaded liposomes, we sought to evaluate how this targeting effects adaptive immune responses. We wanted to confirm that ablating the activity of the FR state would lower the adaptive immune response. However, as the FR state is transient and only exists, by our estimate, for 30 mins, this proved a difficult task. We designed an experiment which would temporarily suppress the FR responses by delivery the protein trafficking inhibitor, BrefA (brefeldin A) we had validated in vitro, with the liposomes – temporarily preventing FRs from signaling via paracrine signaling. We loaded brefA at 1 mg/ml into 200 nm liposomes and observed an approximately 50% loading in both FR-targeted liposomes (FR-TLs) and non targeted liposome (NTL) and validated that these liposomes suppress BMDC activation in vitro (Figure S-20, Figure S-21). After validation, we injected these brefA formulations into mice (100 µg per mouse, i.p.) and then 1 hr later mice were injected i.p with 10 µg of R848 and 100 µg of OVA (free formulation). Controls included liposomes without BrefA (No BrefA), an equivalent free quantity of BrefA (Free), and liposomes with BrefA, but no targeting agent (NTLs). On day 14, mice were boosted using identical conditions. Mice were then sacrificed on day 21 and their anti-OVA titers were analyzed. We observed significant decrease in anti-OVA titers for the FR-TLs compared to NTL (Figure 7A). Furthermore, the FR-TL mice also had decreases in their in CD8/MHCI+ and CD4/MHCII+ cells when compared to untargeted lipo (Figure 7B). However, there was no sigificant change in CD45+ cell populations or any other lymphocyte cell population (Figure S-22). There was a significant decrease in INFγ+ CD4 and CD8 T cells in the ICS analysis in response to the MHCI epitope but not to the MHCII epitope (Figure S-23). This data indicates that by inhibiting the FR population in vivo, the adaptive immune responses can be selectively diminished.

**Figure 7.**
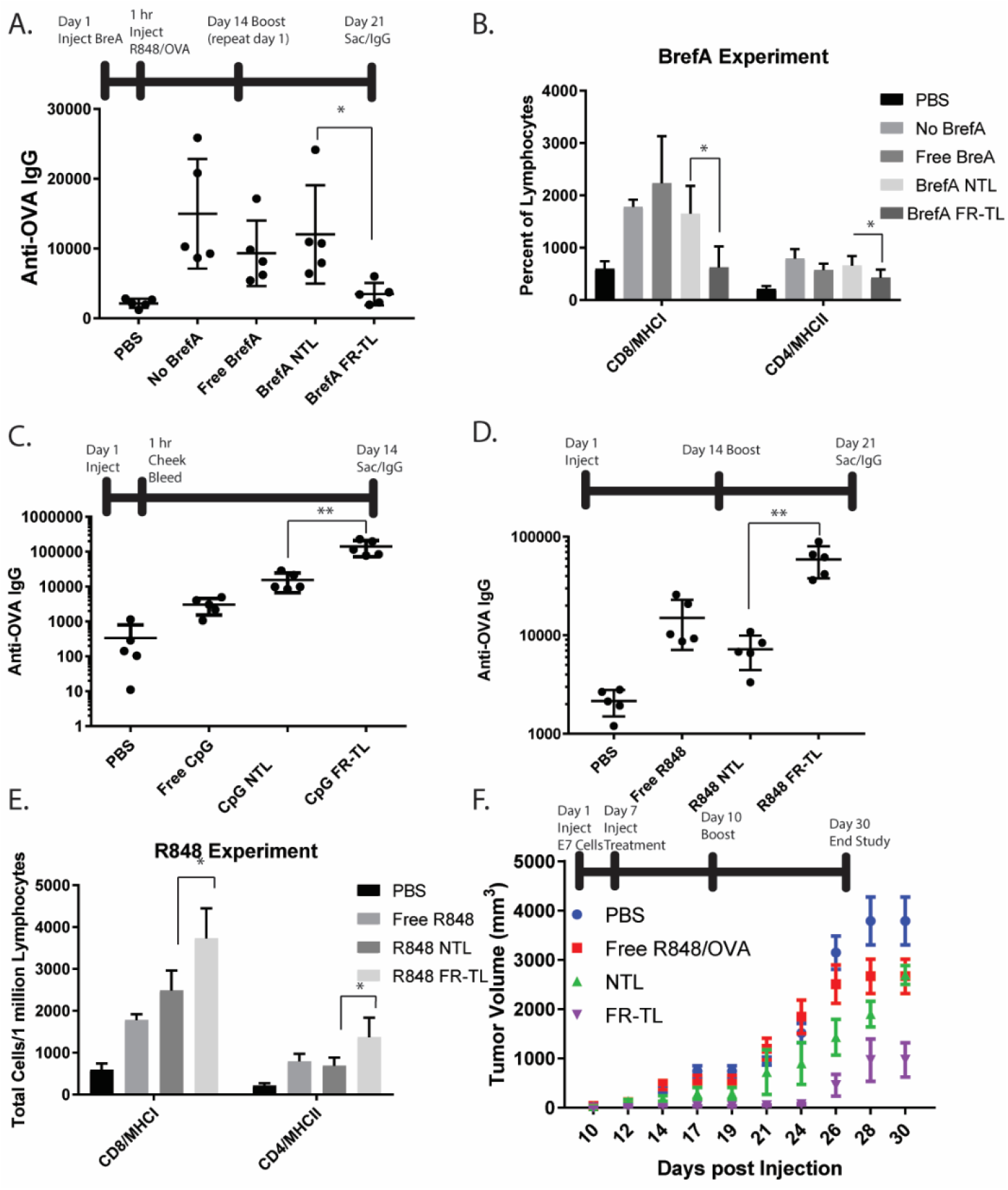
In vivo validation of FR targeting in vaccination models. (A) C57BL/6 mice were injected with Brefeldin A formulations (100 µg per mouse) and then 1 hr later injected with 100 µg OVA and 10 µg CpG on day 1. On day 14 both injections were repeated. Mice were sac’d on day 21 and their IgG titers analyzed. (B) Lymph nodes from experiment in figure 7A were stained for T cell surface markers and with tetramers for the major OVA MHCI epitope (OVA 257-264) and the major MHCII epitope (OVA 323-339) and analyzed via flow. (C)CpG and OVA were loaded into liposomes either targeted with DAP12/Hep (GpG FR-TL) or without (GpG NTL) and injected into C57BL/6 mice and compared to a PBS control or free CpG OVA equivalent (100 µg OVA and 10 µg CpG per mouse). 14 days later anti-OVA IgG titers were measured. (D) R848 (10 µg per mouse) replaced CpG in a repeat of experiment from figure 7A. Mice were injected on day 1 and day 14 and then sac’d and anti-OVA IgG titers were measured on day 21. CD45+ CD3+ CD4+ MHCII tetramer+ and CD45+ CD3+ CD8+ MHCI tetramer+ cells are shown. (E) Aurora analysis similar to part c of lymph nodes of mice from part d. parts a-e, N=5, error bars indicate ±SD and all injections were i.p. (F) E7-OVA tumor model. C57BL/6 mice (N=5) were injected with 2×105 E7-OVA tumor cells in the left flank s.c. on day 1 and tumors measured 3x per week. On day 7 and day 10, mice were injected i.p. with same R848/OVA formulations from part b. Tumors were tracked until day 30. Error bars indicate ±SEM. * indicates p<0.05, ** indicates p<0.01

After demonstrating that FRs played a critical role in vivo, we were intrigued and wondered if the FR-targeted liposomes could be used to enhance conventional responses to adjuvants. We decided to observe the effect of FR targeting on the model antigen ovalbumin (OVA) in three separate experiments. First, we loaded OVA (1 mg/mL) and CpG (a TLR 9 agonist, 200 µg/mL) into FR-TL or NTL (Figure S-19).^29^ We injected these liposomes or an equivalent amount of free CpG/OVA i.p. into C57BL/6 mice (100 µg OVA, 10 µg CpG per mouse). On day 14, there was a nearly 10-fold increase in anti-OVA IgG titers CpG FR-TLs) compared to NTL (Figure 7C). Interestingly, when the mice serum cytokine levels were measured at 1 hr post injection, we did not observe an increase in TNFα levels, indicating that the FR targeting did not increase the systemic cytokine release (Figure S-24).

As the FR state had been responsive to multiple TLRs previously, we explored if the FR-targeted liposomes changed the TLR agonist from CpG to R848, a TLR 7 agonist (Figure S-18).^30^ C57BL/6 mice were injected with R848/OVA formulations on day 0 and 14 and then sacrificed on day 21. The data shows that FR targeting significantly improves anti-OVA IgG levels on day 21 even when changing the TLR agonist and the experimental design (Figure 7D). Furthermore, popliteal and ingunial lymph nodes were removed from both sides of the mouse on day 21, homogenized, stained for various immune cell markers and tetramers to the major MHCI and MHCII epitopes and analyzed via spectral flow cytometry. As shown in figure 7E, targeting FRs significantly increases the number of both MHCI tetramer positive CD8+ T cells and MHCII tetramer positive CD4+ T cells. This increase seems to be a result of an overall increase in CD45+ cells into the lymph nodes and not a disproportionate increase in T cells (Figure S-25A). Furthermore, there is an increase in various T cell subtypes in the FR-targeted group, indicating that FR targeting increases overall cell proliferation instead of a particular cell subtype (Figure S-25B). The spleenocytes from these mice were also analyzed via intracellular staining (ICS) to observe the responses of T cells ex vivo when stimulated by exogenous OVA peptides. There were significant increases in INFγ expressing CD4 and CD8 T cells in responses to the major MHCI OVA peptide and an increase in both INFγ and IL-4 expressing CD4 cell in responses to the major MHCII OVA peptide in the FR targeted group when compared to untargeted liposomes, indicating a broad increase in CD4 and CD8 as well as Th1 and Th2 responses, although the increase in Th1 responses to the MHCI epitope was more dramatic (Figure S-26).

Finally, as we saw increases in many parts of the immune repsonse, we wanted to determine if the antigen-specific responses elicited by FR-targeting could be translated into therapeutic models. For a rudimentary test of this, we used a OVA expressing tumor model (EGF7.OVA) to show that FR targeting alters immune responses in a disease model.^31^ We hypothesized that due to the increase in MHCI CD8 T cell responses from FR targeting, we would observe stronger anti-tumor responses. We injected C57BL/6 mice (n=5) with 2⨯10^5^ E7-OVA cells on day 1 followed by injections of an R848/OVA vaccine formulations or a PBS blank on day 7 and day 10. Tumor volume was measured over a 30 day period during which the mice sacrificed when tumor reached >20 mm in any direction in accordance with our protocol. There was a significant reduction in tumor volume for both liposomal formulation when compared to PBS controls beginning at day 14, but there was a further decrease in tumor volume for the targeted formulation compared to the non-targeted which continued for the remainder of the study (Figure 7F, p<0.05). For example, on day 24, the FR-TL group had an average tumor volume of 70±37 mm^3^, while all other groups had an average tumor volume >800 mm^3^. On day 30, all mice except for the FR-TL group were sacrificed due to tumor size (Figure S-27). Taken together, these experiments indicate that FR targeting does improve overall immune reponses in vivo by increasing CD8 T cell and IgG responses.

## Conclusion

In this study, we started with the observation of a statistical anomaly in particle uptake and followed that to provide in vitro and in vivo evidence of a transcriptionally distinct cellular state of dendritic cells belonging predominantly to the cDC2 subset. This state, which we refer to as First Responders, is identified as having increased sensitivity to TLR signaling and disseminates global APC activation via paracrine signaling. We present evidence that this state is temporally controlled, with cells maintaining a high activation state for hours before returning to a “non-FR” state. Furthermore, we observed that activation of the FR cells is a large component of adaptive immune responses both for adoptive transfer experiments and for loss of function experiments in vivo. We also have identified several proteins that are transcriptionally upregulated when a cell is in the FR state and note two, DAP12 and PRG2, that can be effectively targeted both in vitro and in vivo to identify FRs from other dendritic cells. This targeting showed promise to improve model vaccination efficacy when used in vivo vaccination with OVA.

While the identification of a potential FR state has a wide impact on both our scientific understanding of how innate immune cells respond to pathogens and a wide range of therapeutic potential for FR targeting, we wish to acknowledge several experimental elements in this study that will require further expansion before FR state can be fully vetted. Our kinetic analysis of FR activation was limited; due to the restriction of FRs in our early experiments requiring uptake of a micro-particle for analysis. A more in-depth study will need to rely upon proteins exclusively expressed in FRs and identify them in their naïve state. While we have identified upregulated surface proteins in our mRNA analysis, our initial isolation protocols were limited by the ligands both for in vivo and in vitro analysis. Heparin can have varying immunological effects and, due its heterogeneous structure and low binding affinity to PRG2, it may have other effects that are difficult to decouple from FR targeting.^32^ Also, vaccination with targeting peptides (such as DAP12 peptide) has limited therapeutic applications due to the potential of developing immune responses to the peptide of interest. Furthermore, we do not know the functional importance PRG2 and DAP12 have on the FR state. Further research might help identify proteins more directly involved in FR state regulation and control and provide context on previous research on DAP12 and PRG2. For example, DAP12 is highly associated with NK cells and glial cells and associates with another protein TREM2, which recently has been implicated as an immune regulator in brain cancer.^33^ There may be a connection between TREM, DAP12 and immune regulation that will require further research. Additionally, a more thorough analysis of the kinetics of cytokine release in FR would provide valuable insights into which cytokines are critical for paracrine signaling. Finally, our work is exclusively conducted in mice and it remains to be seen if FR populations exist in human DCs. Despite these limitations, we present this data as an important first evaluation of FRs so that others might explore their role in innate and adaptive immune function.

One large remaining question that we do not yet have an answer for is why do DCs appear to cycle through these activation states? For now, we can only conjecture that this is due to the large metabolic strain placed on DCs when activated and controlling the activation of neighboring DCs, requiring DCs to “share the load”. DCs release many cytokines by exosome, which require them to synthesize them prior to activation, in order to have rapid responses to pathogens.^34^ We hypothesize that DCs might cycle through this state of increased readiness in order to reduce overall metabolic load on individual cells, while relying on the high sensitivity of immune cells to cytokine signaling. This sort of cooperativity has a precedent in other antigen presenting cells; macrophages exist in at least two distinct metabolic states, M1 and less active M2, which have been suggested to divide up the metabolic strain between M1 and M2.^35^ Furthermore, previous work has shown that DCs which have been previously activated can transmit their activation to naïve cells, suggesting a similar paracrine signaling cascade.^36^ While much further research will be required to confirm this, this work suggests a new approach to studying DCs activation and propagation of immune signals. Identifying and communicating with FR states has most clear implications for therapies where targeted immune activation is desired including vaccines and immunotherapies, but perhaps FRs also play a role in other disease in which innate immunity is implicated. The identification of FRs provides a new insight into how immune responses are modulated by innate immune cells states and targeting FRs can potentially help improve therapeutics where innate immune activation is required.

## Materials and Methods

See Supplementary Information for details on materials and methods.

## Acknowledgements

We would like to thank Dr. Anne Sperling, Dr. Jacques Banchereau, Dr. Peter Savage and Nicolas Chevrier for their feedback on our manuscript and experimental plan.

## Author contributions

PD planned and performed the majority of experiments, prepared figures, wrote the majority of the manuscript, edited the manuscript and synthesized FR targeting liposomes. BS synthesized TLR conjugated MP, performed experiments, analyzed data and wrote sections of the paper. RS developed TLR conjugated MP synthesis, designed experiments and edited the paper. AEK conceived the idea for the study, designed experiments, wrote sections of the paper and edited the paper.

## Materials & Correspondence

Correspondence and material requests should be addressed to Dr. Aaron Esser-Kahn

