## Supplemental Figures for "Identification, Characterization, and Targeting of a Rare and Temporal Dendritic Cell State that Facilitates Adaptive Immune Responses"

### Slide 1
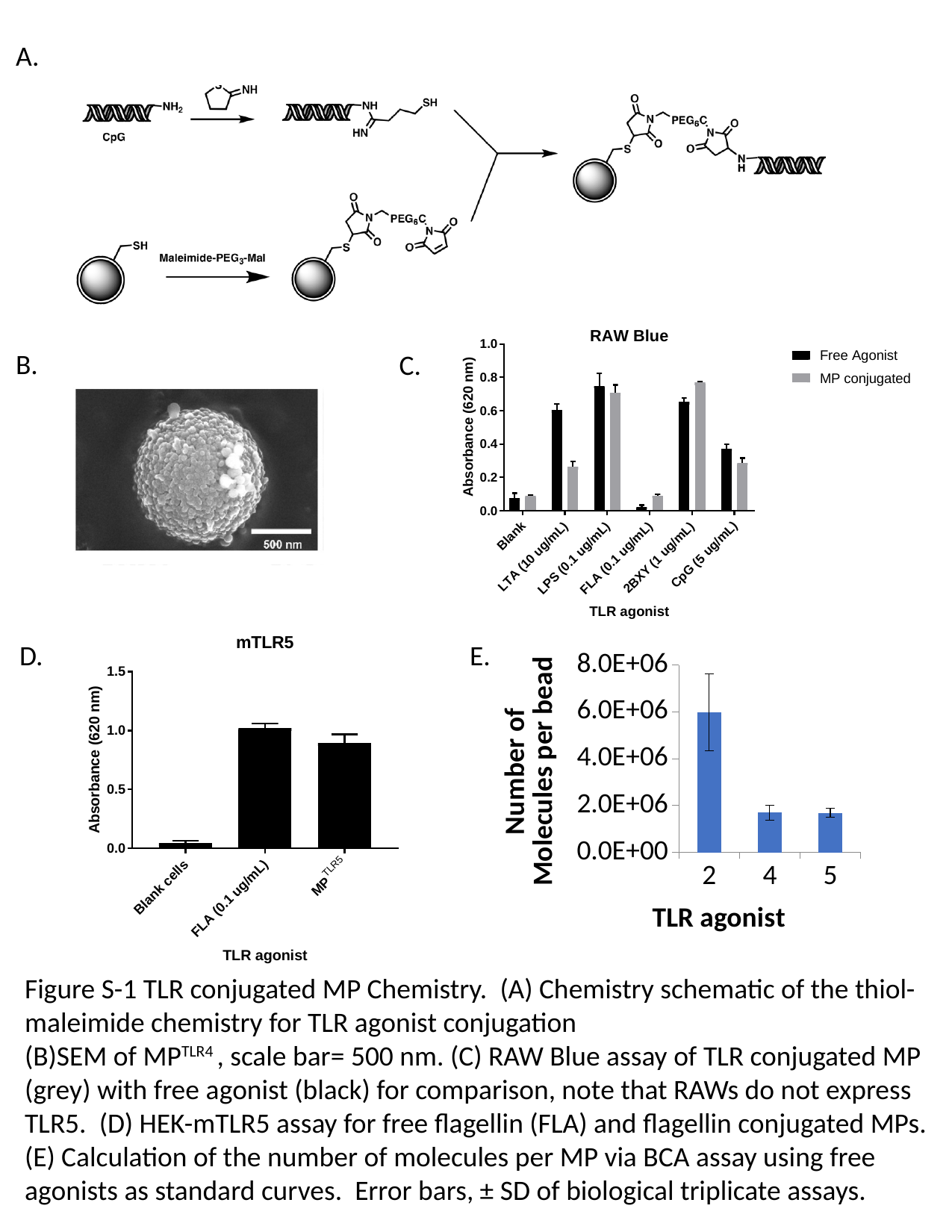

A.
B.
C.
D.
E.
###### Chart
| Category | |
|---|---|
| 2 | 5975735.294117646 |
| 4 | 1693626.666666667 |
| 5 | 1693125.0000000005 |Figure S-1 TLR conjugated MP Chemistry. (A) Chemistry schematic of the thiol-maleimide chemistry for TLR agonist conjugation
(B)SEM of MP^TLR4^ , scale bar= 500 nm. (C) RAW Blue assay of TLR conjugated MP (grey) with free agonist (black) for comparison, note that RAWs do not express TLR5. (D) HEK-mTLR5 assay for free flagellin (FLA) and flagellin conjugated MPs. (E) Calculation of the number of molecules per MP via BCA assay using free agonists as standard curves. Error bars, ± SD of biological triplicate assays.

### Slide 2
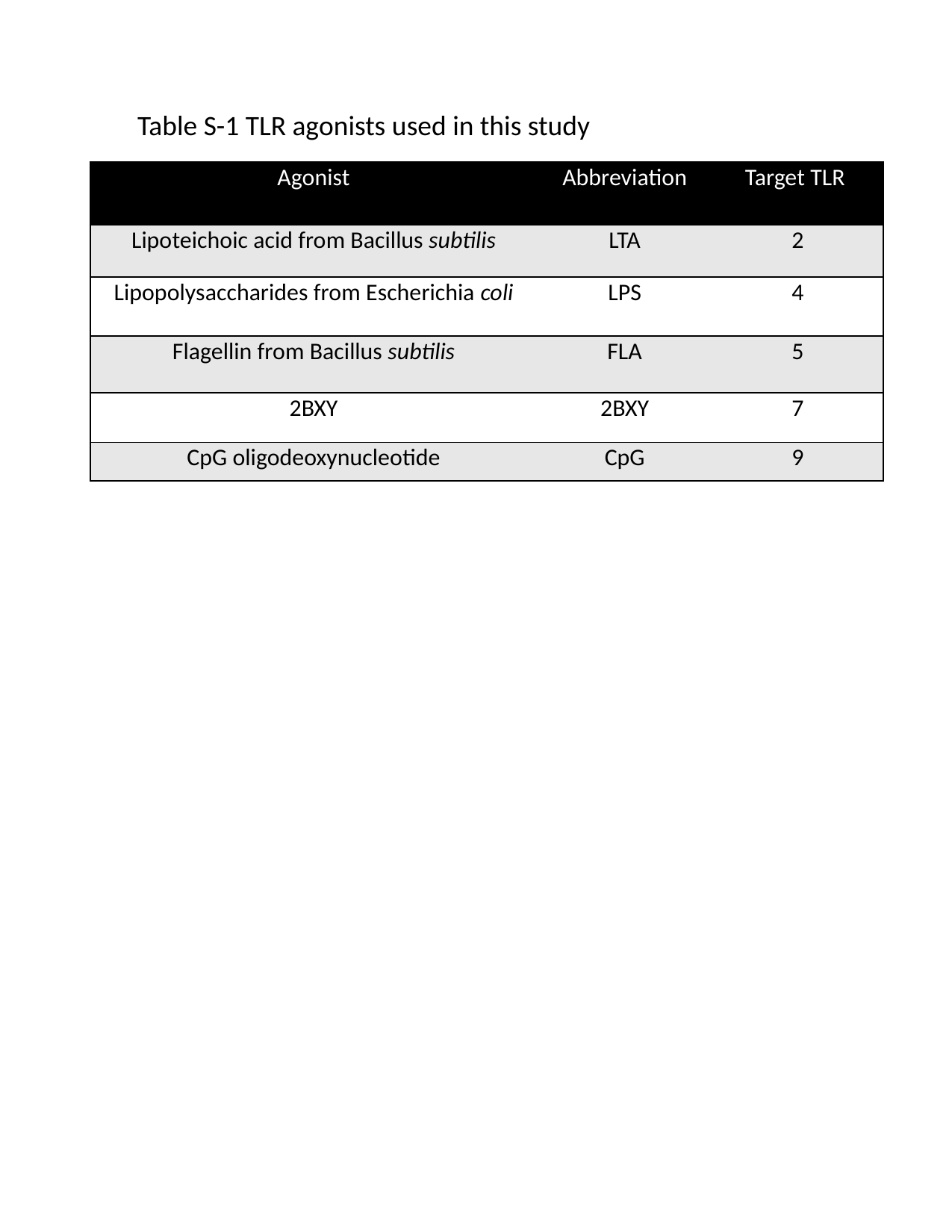

Table S-1 TLR agonists used in this study
| Agonist | Abbreviation | Target TLR |
| --- | --- | --- |
| Lipoteichoic acid from Bacillus subtilis | LTA | 2 |
| Lipopolysaccharides from Escherichia coli | LPS | 4 |
| Flagellin from Bacillus subtilis | FLA | 5 |
| 2BXY | 2BXY | 7 |
| CpG oligodeoxynucleotide | CpG | 9 |

### Slide 3
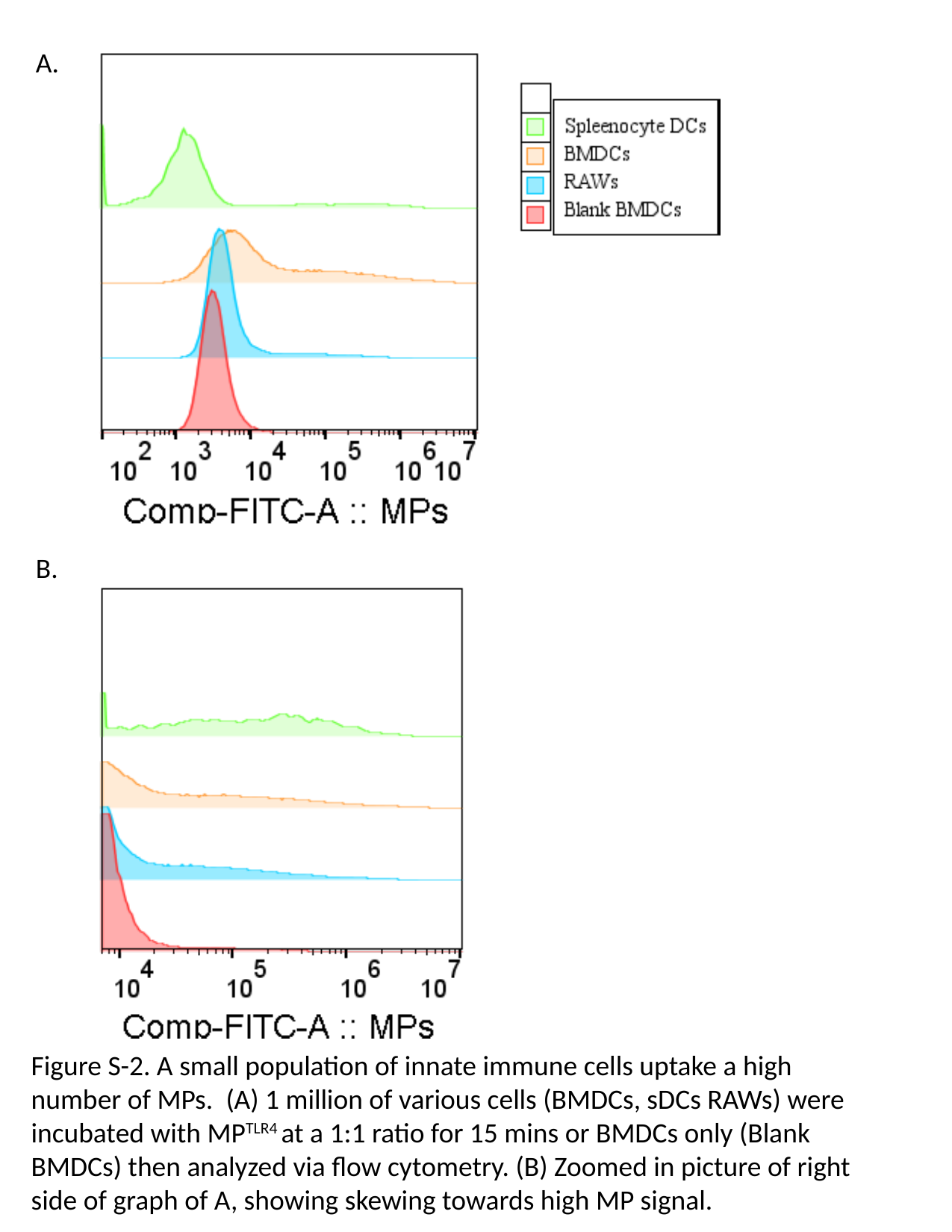

A.
B.
Figure S-2. A small population of innate immune cells uptake a high number of MPs. (A) 1 million of various cells (BMDCs, sDCs RAWs) were incubated with MP^TLR4^ at a 1:1 ratio for 15 mins or BMDCs only (Blank BMDCs) then analyzed via flow cytometry. (B) Zoomed in picture of right side of graph of A, showing skewing towards high MP signal.

### Slide 4
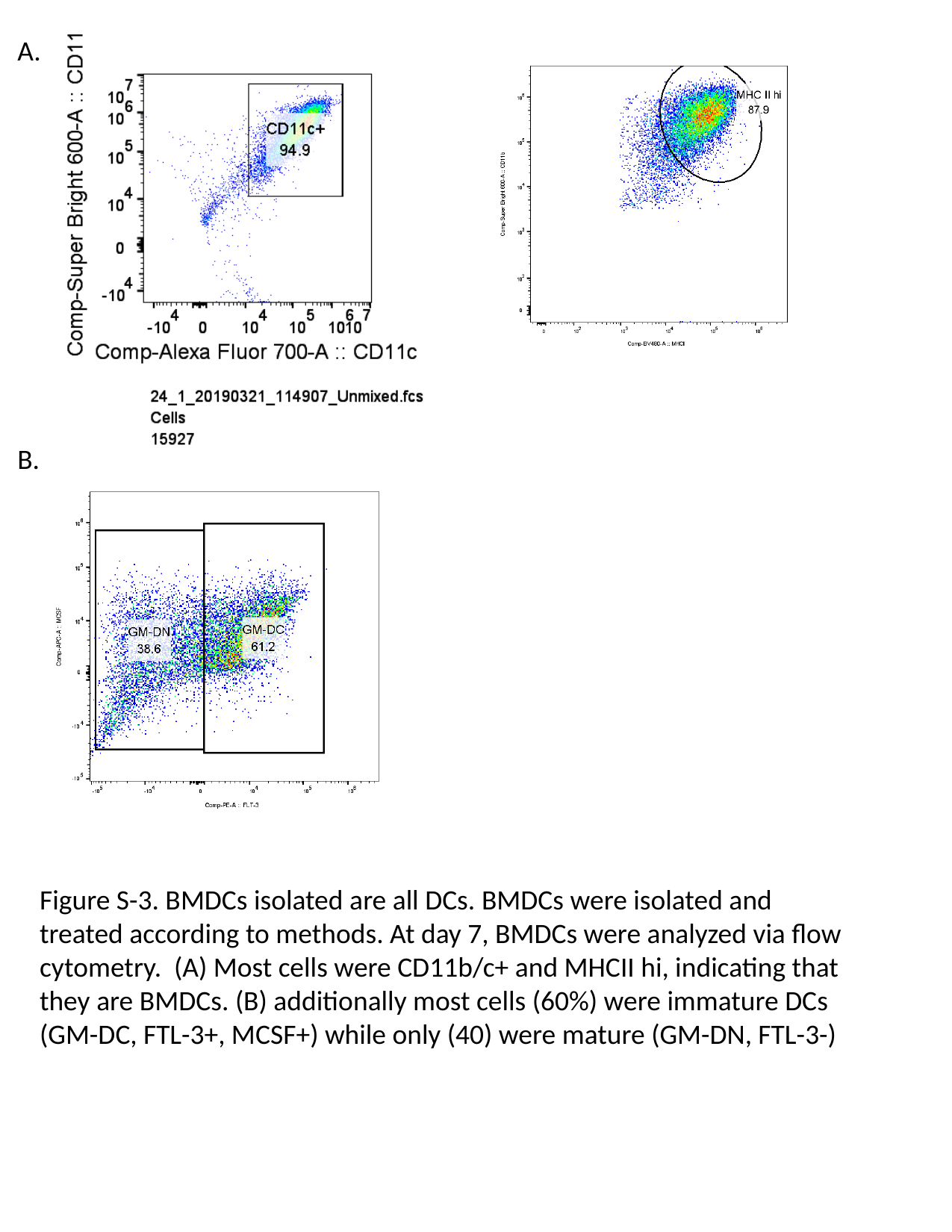

A.
B.
Figure S-3. BMDCs isolated are all DCs. BMDCs were isolated and treated according to methods. At day 7, BMDCs were analyzed via flow cytometry. (A) Most cells were CD11b/c+ and MHCII hi, indicating that they are BMDCs. (B) additionally most cells (60%) were immature DCs (GM-DC, FTL-3+, MCSF+) while only (40) were mature (GM-DN, FTL-3-)

### Slide 5
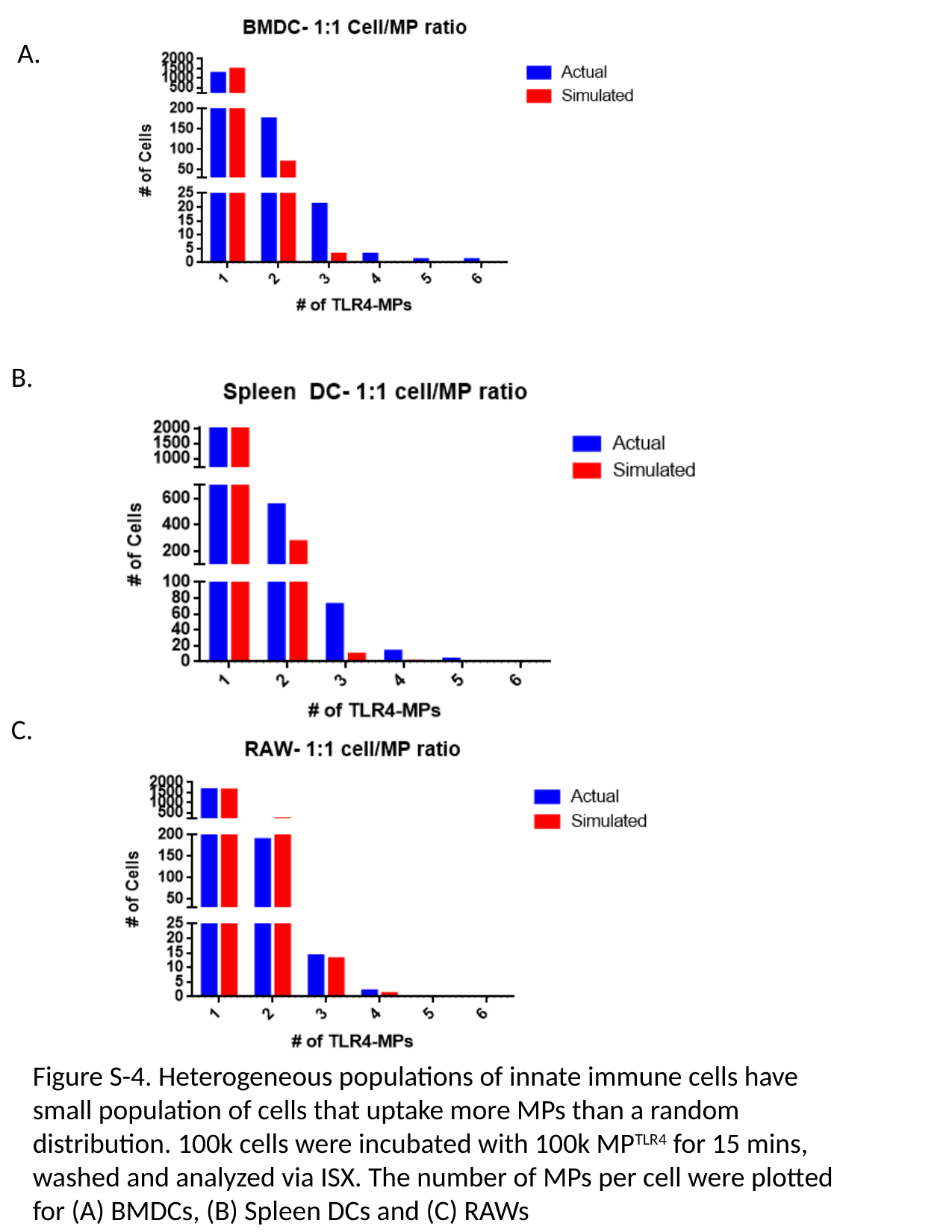

A.
B.
C.
Figure S-4. Heterogeneous populations of innate immune cells have small population of cells that uptake more MPs than a random distribution. 100k cells were incubated with 100k MP^TLR4^ for 15 mins, washed and analyzed via ISX. The number of MPs per cell were plotted for (A) BMDCs, (B) Spleen DCs and (C) RAWs

### Slide 6
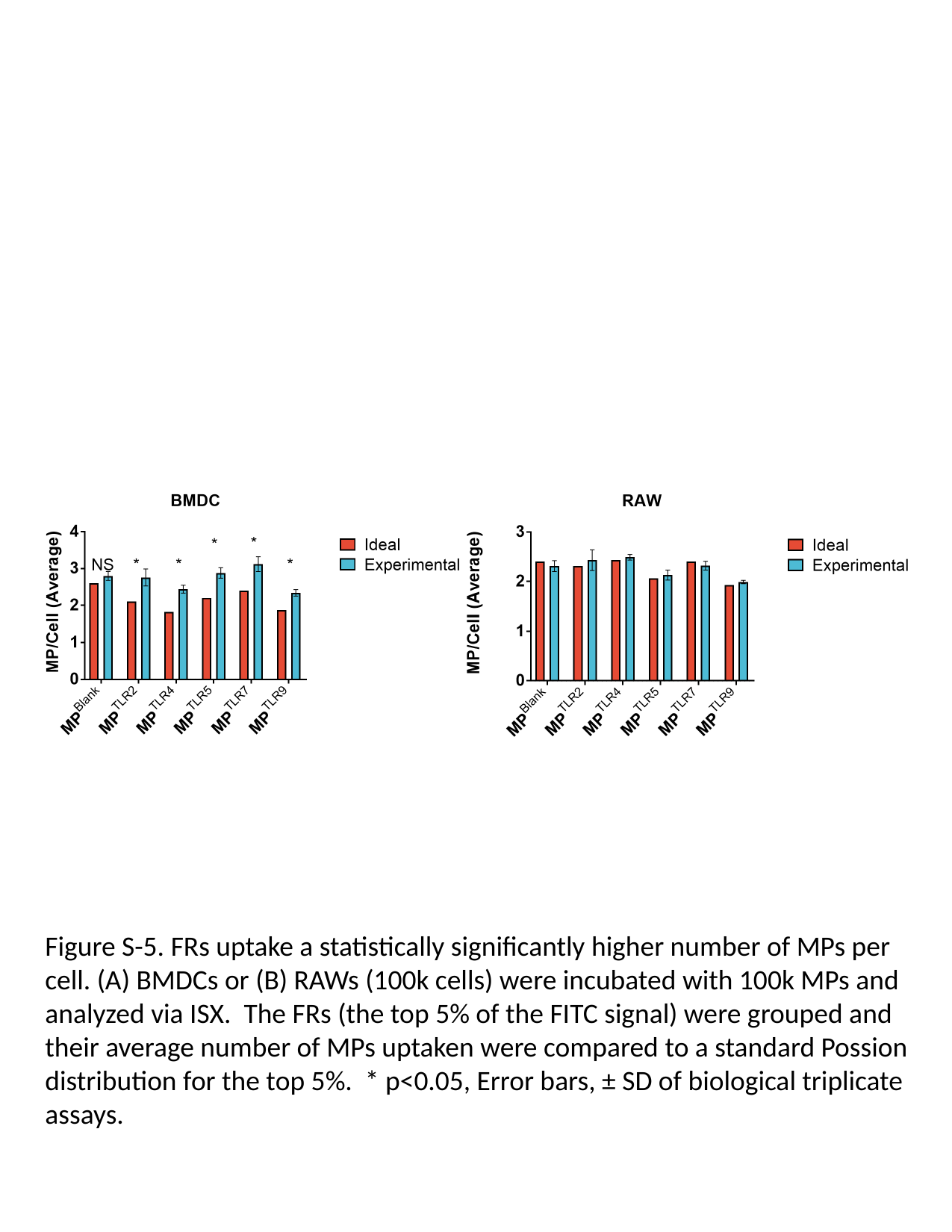

Figure S-5. FRs uptake a statistically significantly higher number of MPs per cell. (A) BMDCs or (B) RAWs (100k cells) were incubated with 100k MPs and analyzed via ISX. The FRs (the top 5% of the FITC signal) were grouped and their average number of MPs uptaken were compared to a standard Possion distribution for the top 5%. * p<0.05, Error bars, ± SD of biological triplicate assays.

### Slide 7
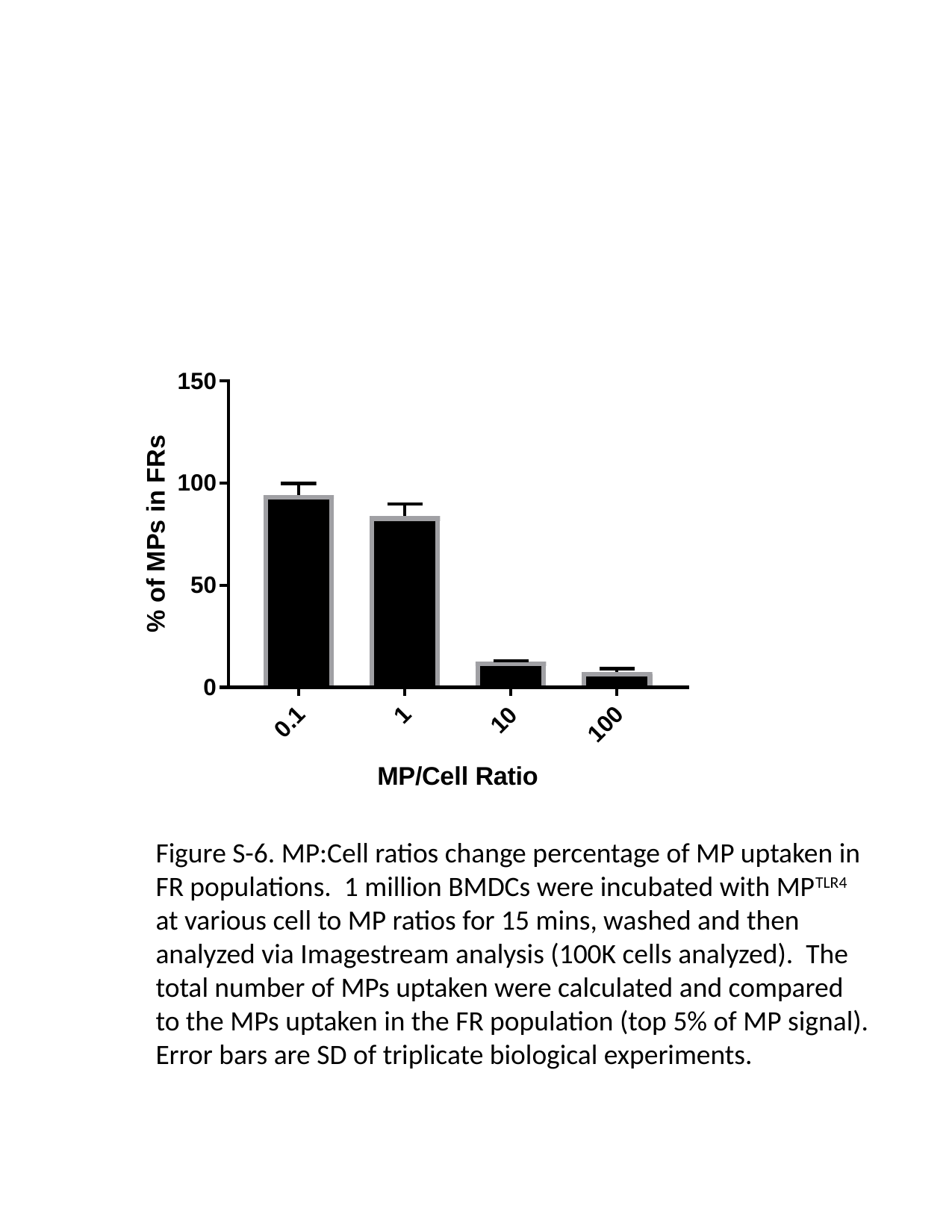

Figure S-6. MP:Cell ratios change percentage of MP uptaken in FR populations. 1 million BMDCs were incubated with MP^TLR4^ at various cell to MP ratios for 15 mins, washed and then analyzed via Imagestream analysis (100K cells analyzed). The total number of MPs uptaken were calculated and compared to the MPs uptaken in the FR population (top 5% of MP signal). Error bars are SD of triplicate biological experiments.

### Slide 8
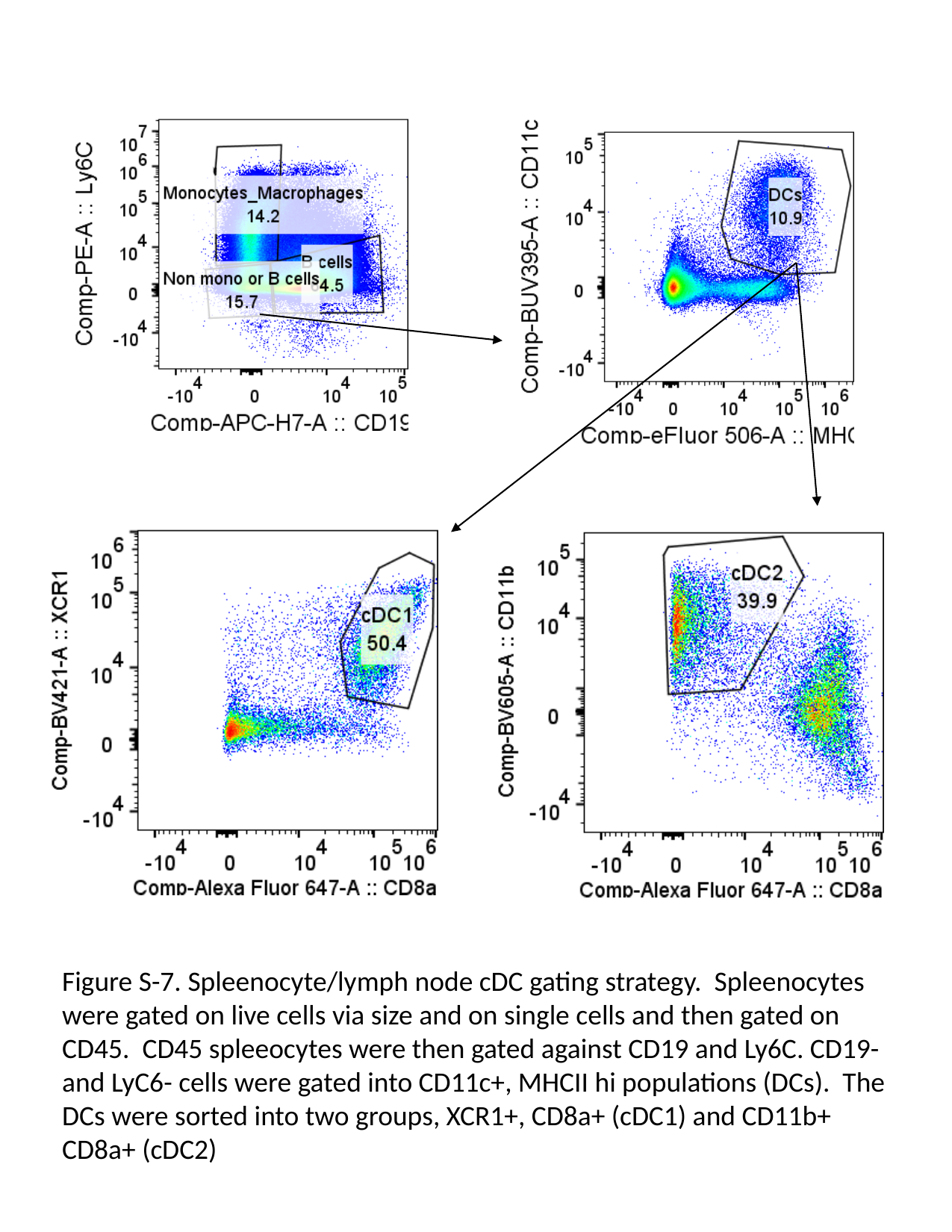

Figure S-7. Spleenocyte/lymph node cDC gating strategy. Spleenocytes were gated on live cells via size and on single cells and then gated on CD45. CD45 spleeocytes were then gated against CD19 and Ly6C. CD19- and LyC6- cells were gated into CD11c+, MHCII hi populations (DCs). The DCs were sorted into two groups, XCR1+, CD8a+ (cDC1) and CD11b+ CD8a+ (cDC2)

### Slide 9
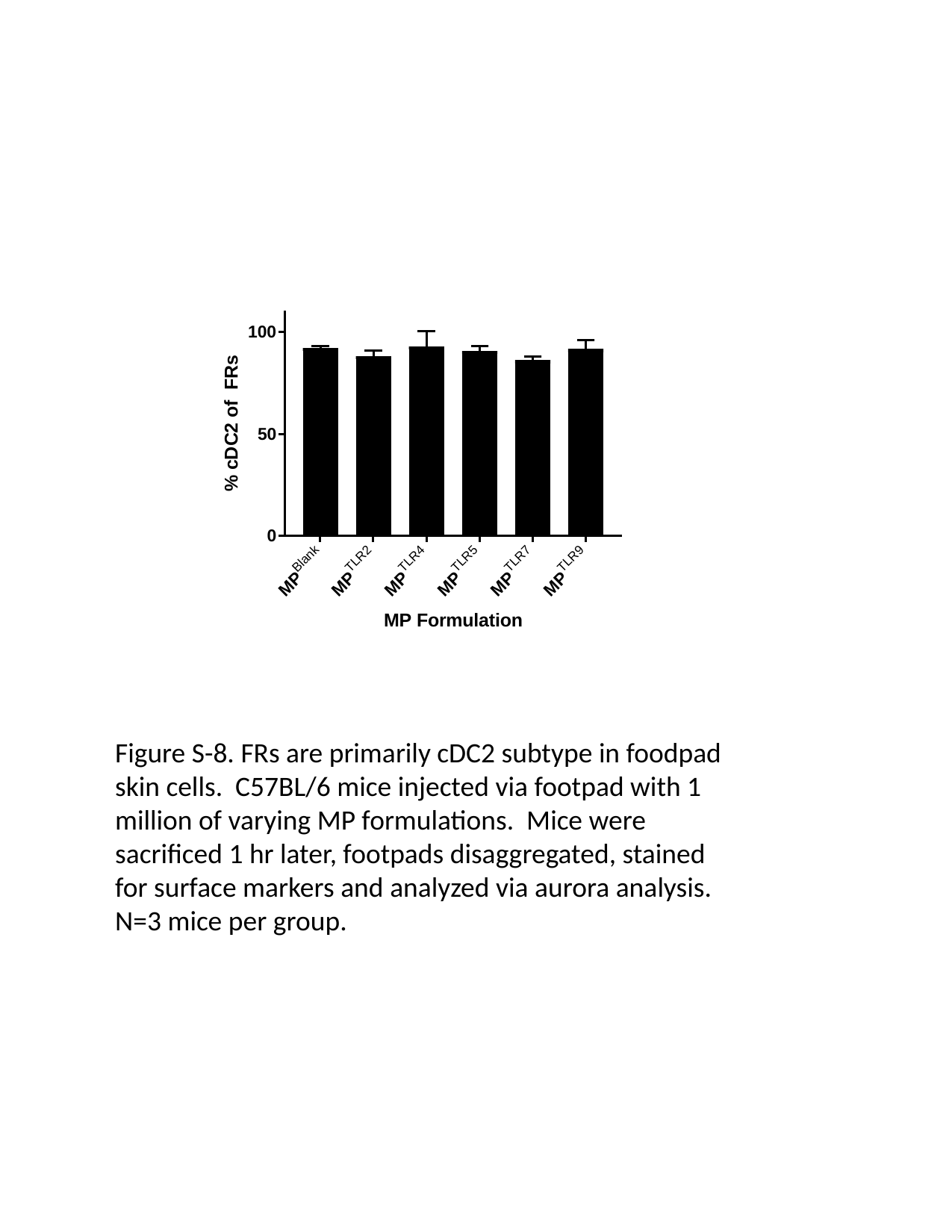

Figure S-8. FRs are primarily cDC2 subtype in foodpad skin cells. C57BL/6 mice injected via footpad with 1 million of varying MP formulations. Mice were sacrificed 1 hr later, footpads disaggregated, stained for surface markers and analyzed via aurora analysis. N=3 mice per group.

### Slide 10
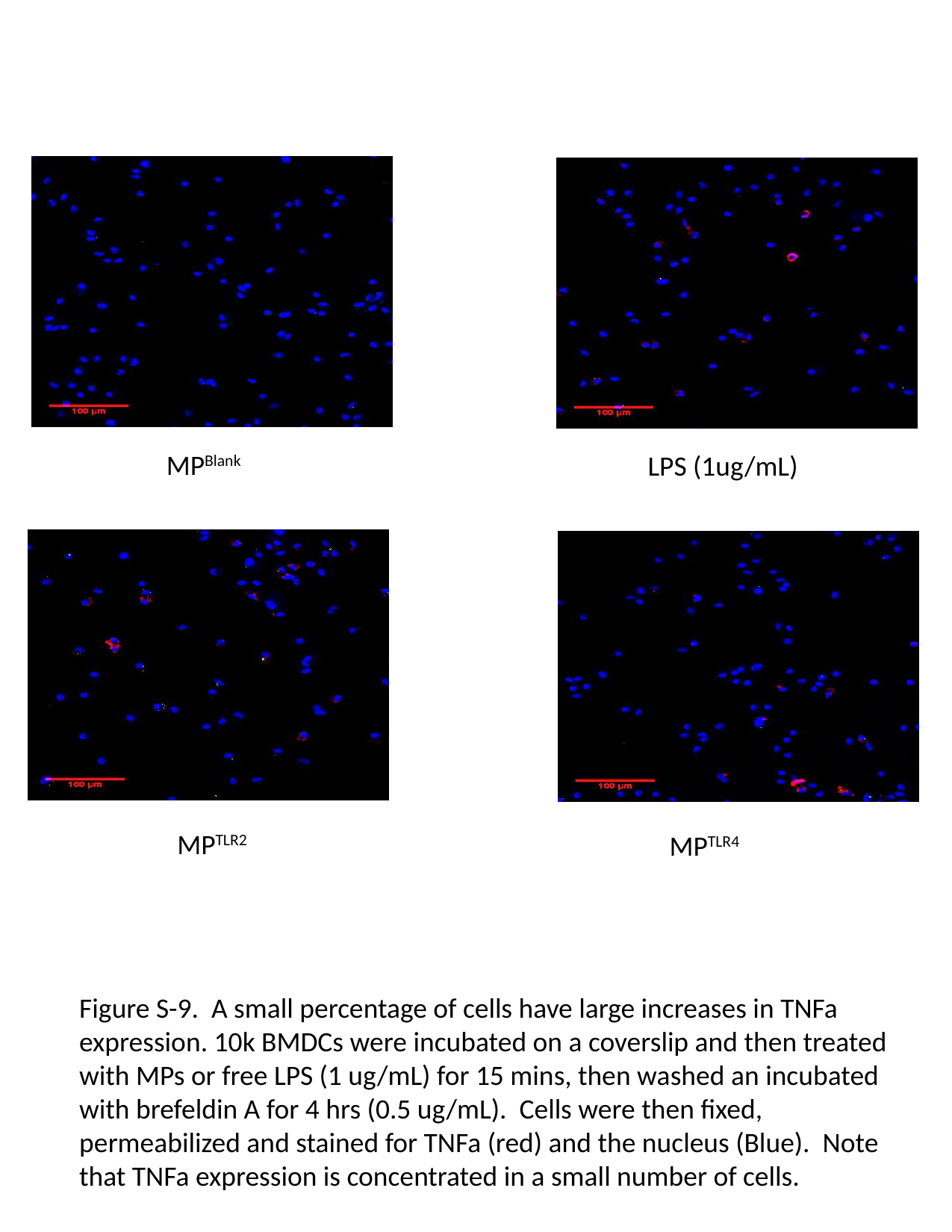

MP^Blank^
LPS (1ug/mL)
MP^TLR2^
MP^TLR4^
Figure S-9. A small percentage of cells have large increases in TNFa expression. 10k BMDCs were incubated on a coverslip and then treated with MPs or free LPS (1 ug/mL) for 15 mins, then washed an incubated with brefeldin A for 4 hrs (0.5 ug/mL). Cells were then fixed, permeabilized and stained for TNFa (red) and the nucleus (Blue). Note that TNFa expression is concentrated in a small number of cells.

### Slide 11
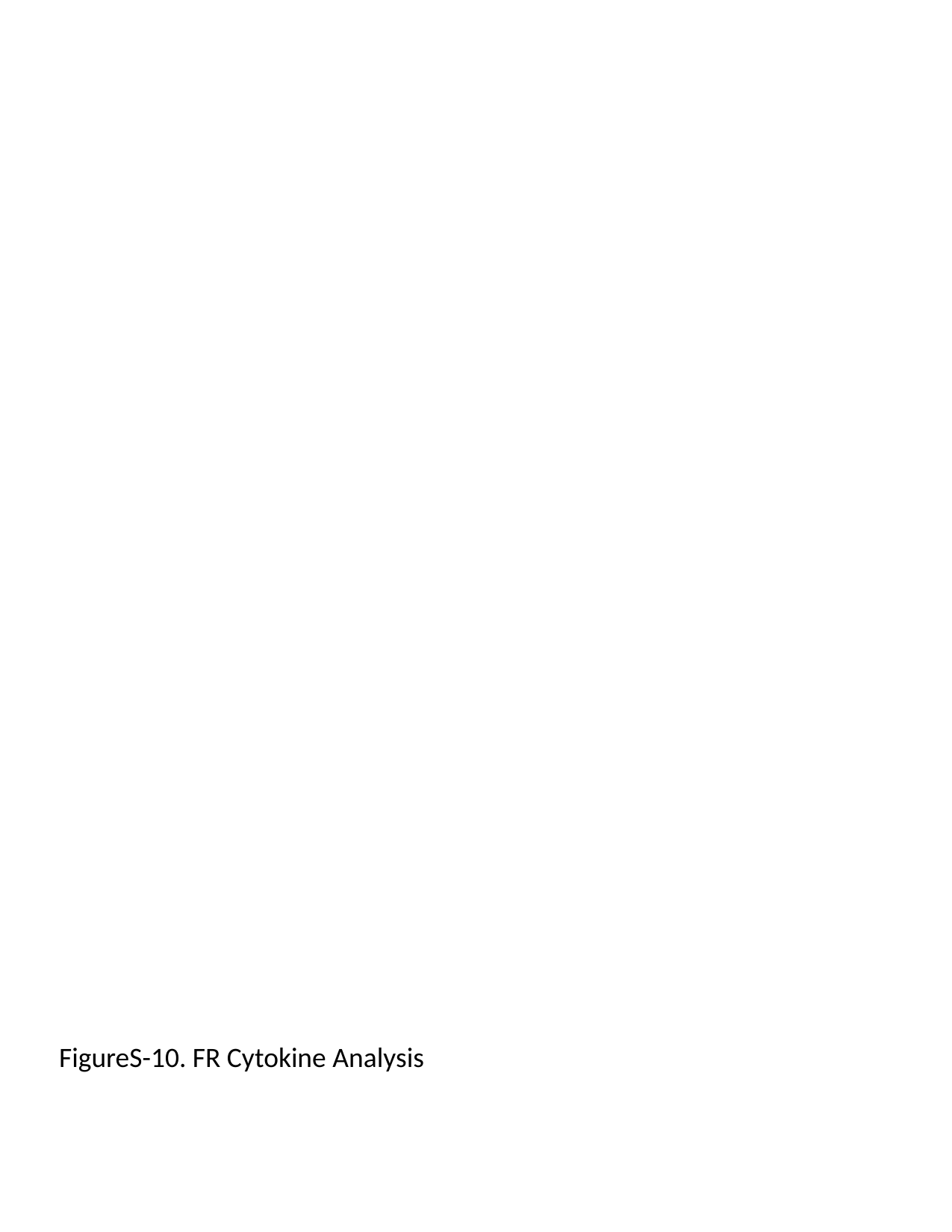

FigureS-10. FR Cytokine Analysis

### Slide 12
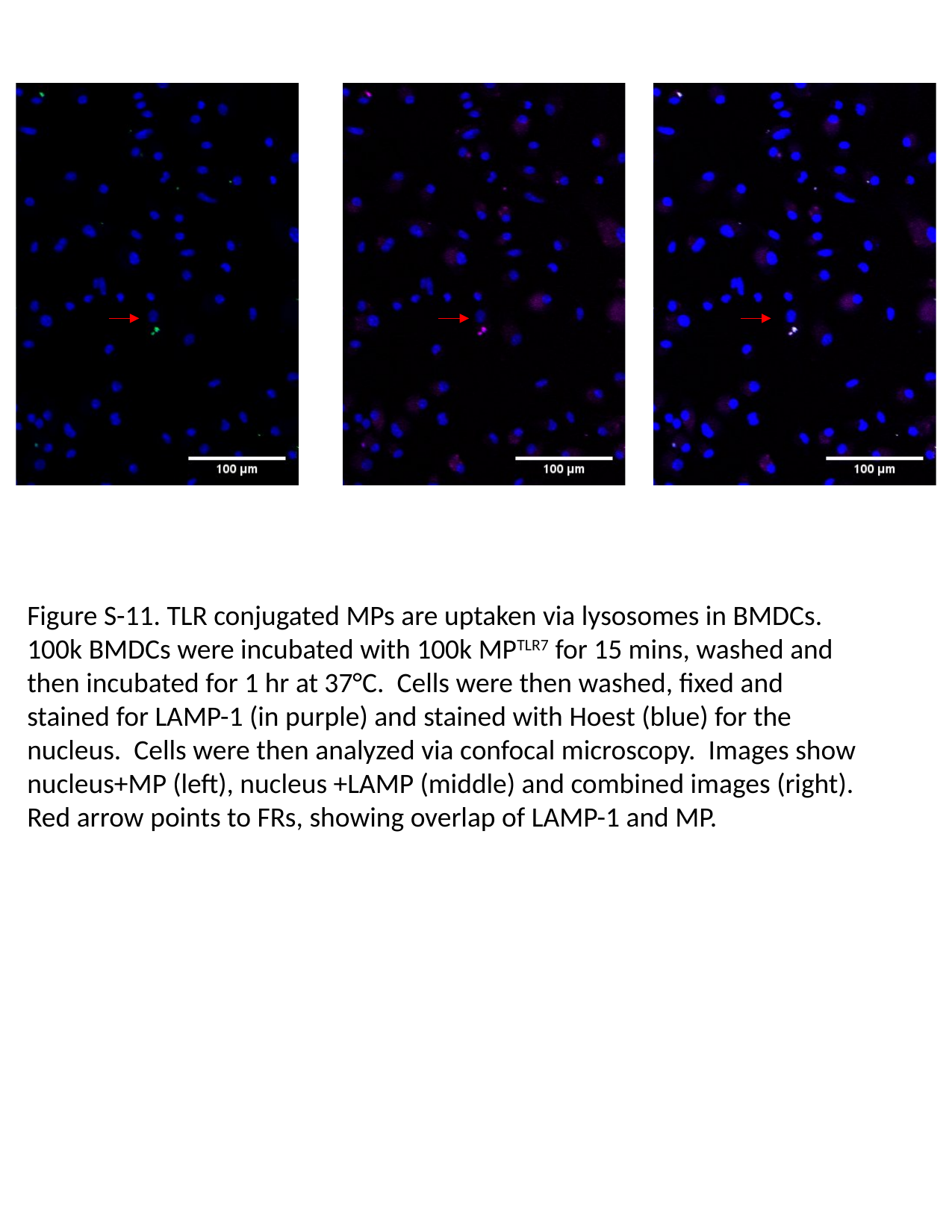

Figure S-11. TLR conjugated MPs are uptaken via lysosomes in BMDCs. 100k BMDCs were incubated with 100k MP^TLR7^ for 15 mins, washed and then incubated for 1 hr at 37°C. Cells were then washed, fixed and stained for LAMP-1 (in purple) and stained with Hoest (blue) for the nucleus. Cells were then analyzed via confocal microscopy. Images show nucleus+MP (left), nucleus +LAMP (middle) and combined images (right). Red arrow points to FRs, showing overlap of LAMP-1 and MP.

### Slide 13
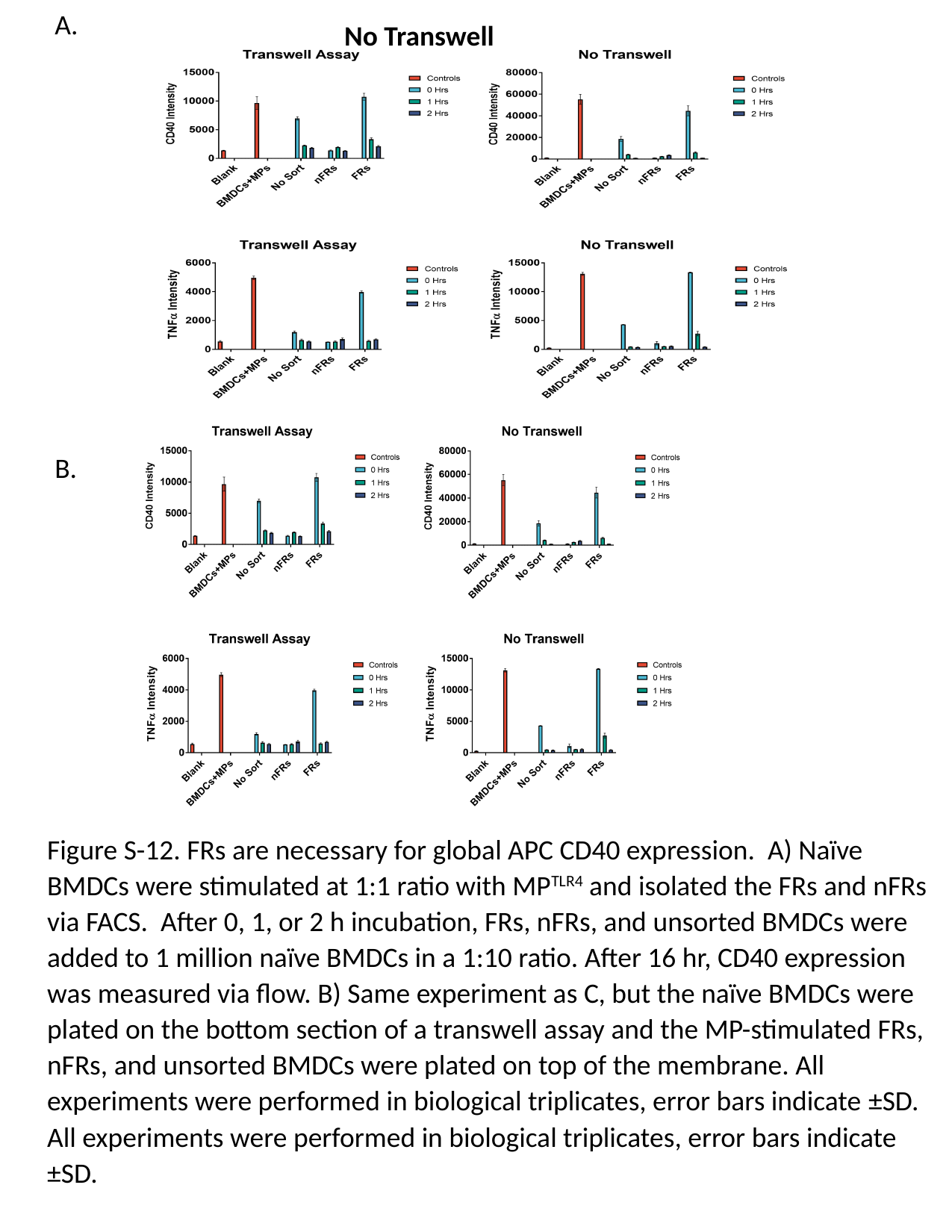

A.
No Transwell
B.
Figure S-12. FRs are necessary for global APC CD40 expression. A) Naïve BMDCs were stimulated at 1:1 ratio with MP^TLR4^ and isolated the FRs and nFRs via FACS. After 0, 1, or 2 h incubation, FRs, nFRs, and unsorted BMDCs were added to 1 million naïve BMDCs in a 1:10 ratio. After 16 hr, CD40 expression was measured via flow. B) Same experiment as C, but the naïve BMDCs were plated on the bottom section of a transwell assay and the MP-stimulated FRs, nFRs, and unsorted BMDCs were plated on top of the membrane. All experiments were performed in biological triplicates, error bars indicate ±SD. All experiments were performed in biological triplicates, error bars indicate ±SD.

### Slide 14
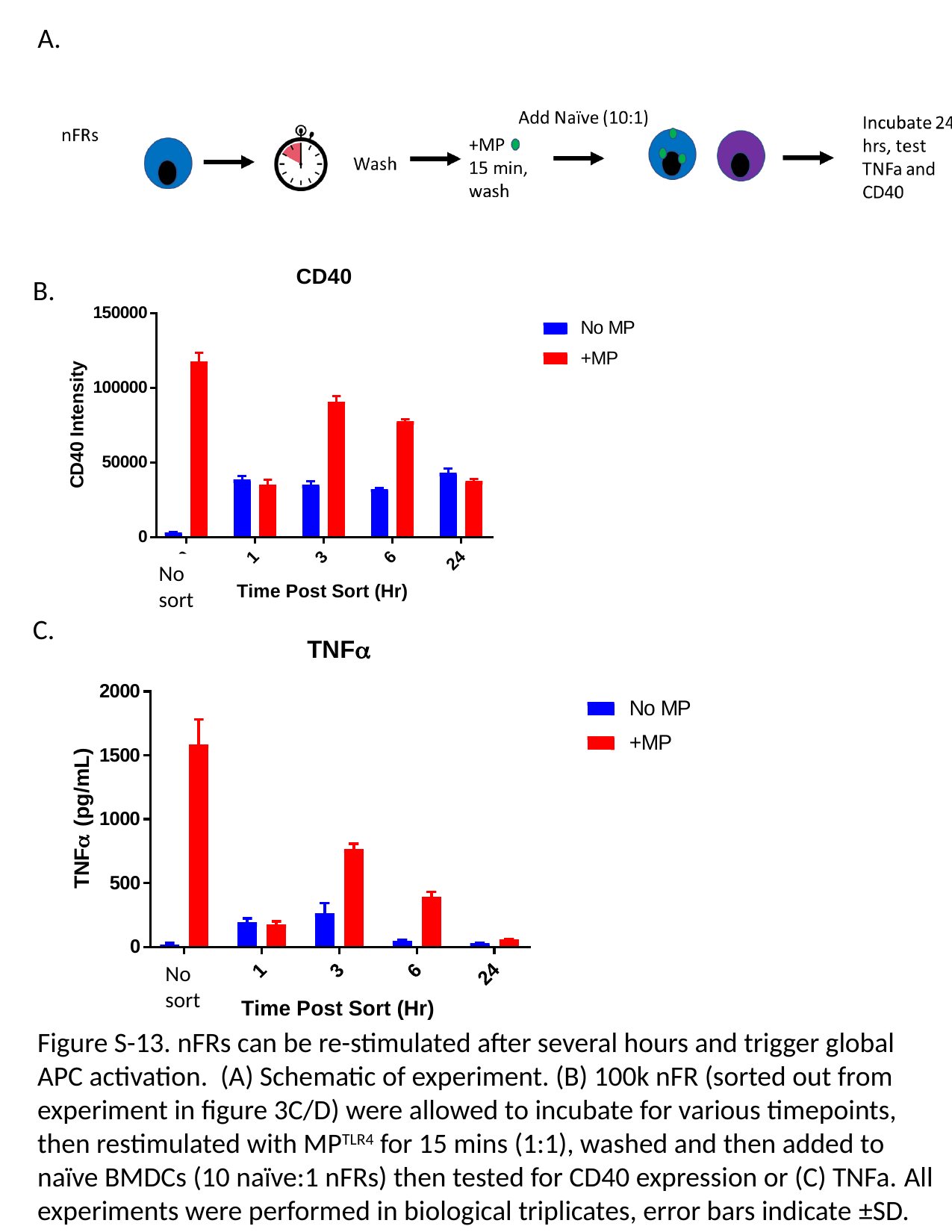

A.
No sort
B.
C.
No sort
Figure S-13. nFRs can be re-stimulated after several hours and trigger global APC activation. (A) Schematic of experiment. (B) 100k nFR (sorted out from experiment in figure 3C/D) were allowed to incubate for various timepoints, then restimulated with MP^TLR4^ for 15 mins (1:1), washed and then added to naïve BMDCs (10 naïve:1 nFRs) then tested for CD40 expression or (C) TNFa. All experiments were performed in biological triplicates, error bars indicate ±SD.

### Slide 15
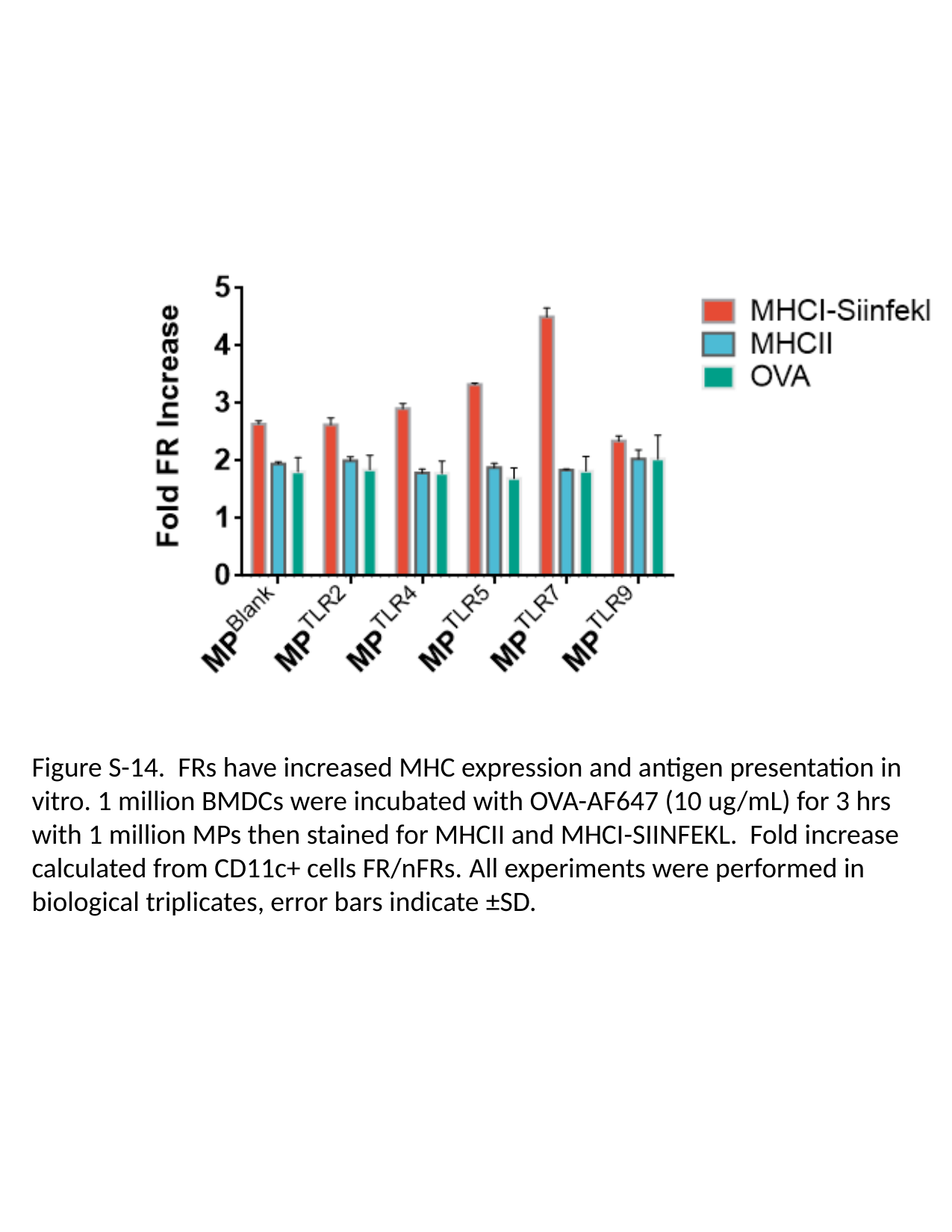

Figure S-14. FRs have increased MHC expression and antigen presentation in vitro. 1 million BMDCs were incubated with OVA-AF647 (10 ug/mL) for 3 hrs with 1 million MPs then stained for MHCII and MHCI-SIINFEKL. Fold increase calculated from CD11c+ cells FR/nFRs. All experiments were performed in biological triplicates, error bars indicate ±SD.

### Slide 16
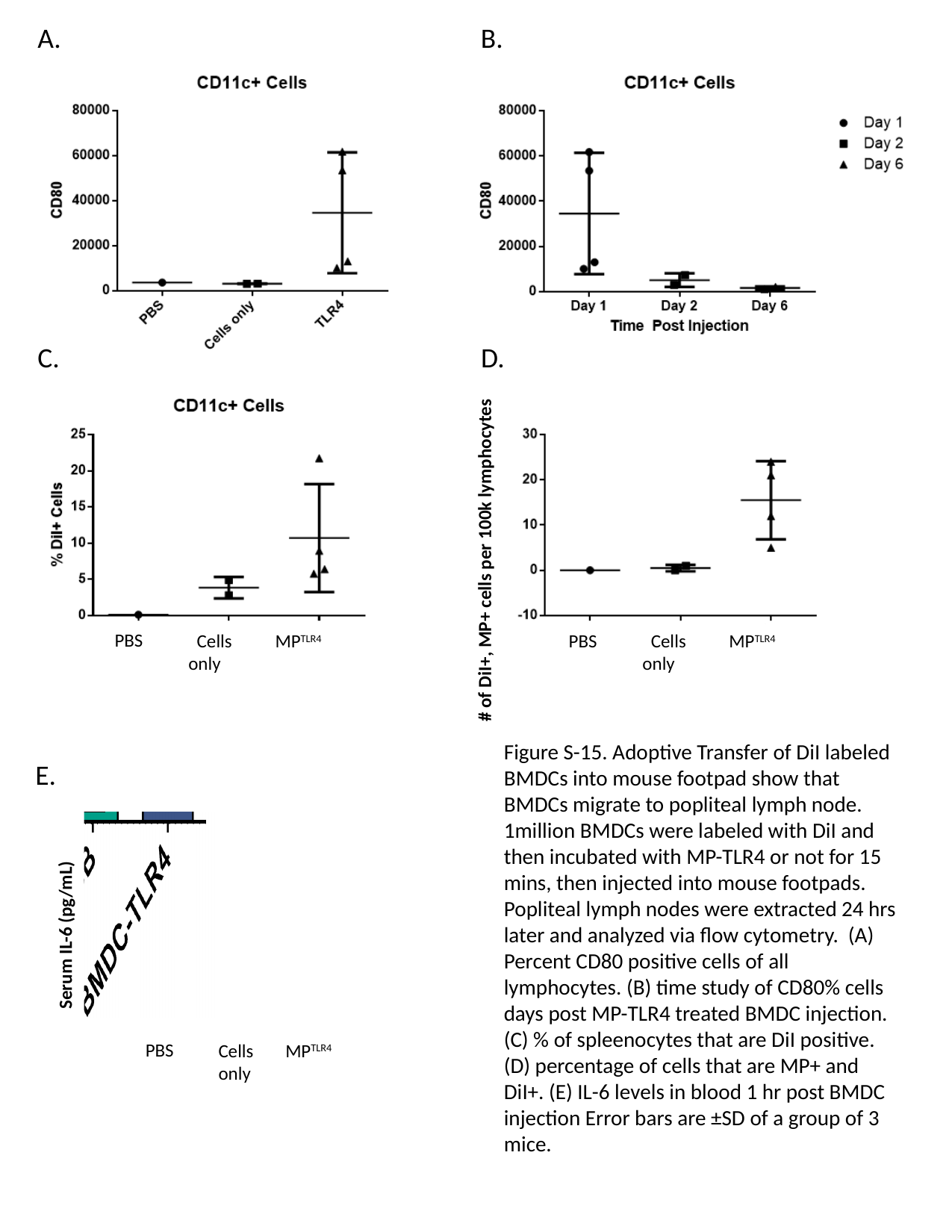

A.
B.
C.
D.
#### of DiI+, MP+ cells per 100k lymphocytes
PBS
 Cells only
MP^TLR4^
PBS
 Cells only
MP^TLR4^
Figure S-15. Adoptive Transfer of DiI labeled BMDCs into mouse footpad show that BMDCs migrate to popliteal lymph node. 1million BMDCs were labeled with DiI and then incubated with MP-TLR4 or not for 15 mins, then injected into mouse footpads. Popliteal lymph nodes were extracted 24 hrs later and analyzed via flow cytometry. (A) Percent CD80 positive cells of all lymphocytes. (B) time study of CD80% cells days post MP-TLR4 treated BMDC injection. (C) % of spleenocytes that are DiI positive. (D) percentage of cells that are MP+ and DiI+. (E) IL-6 levels in blood 1 hr post BMDC injection Error bars are ±SD of a group of 3 mice.
E.
 Serum IL-6 (pg/mL)
PBS
Cells only
MP^TLR4^

### Slide 17
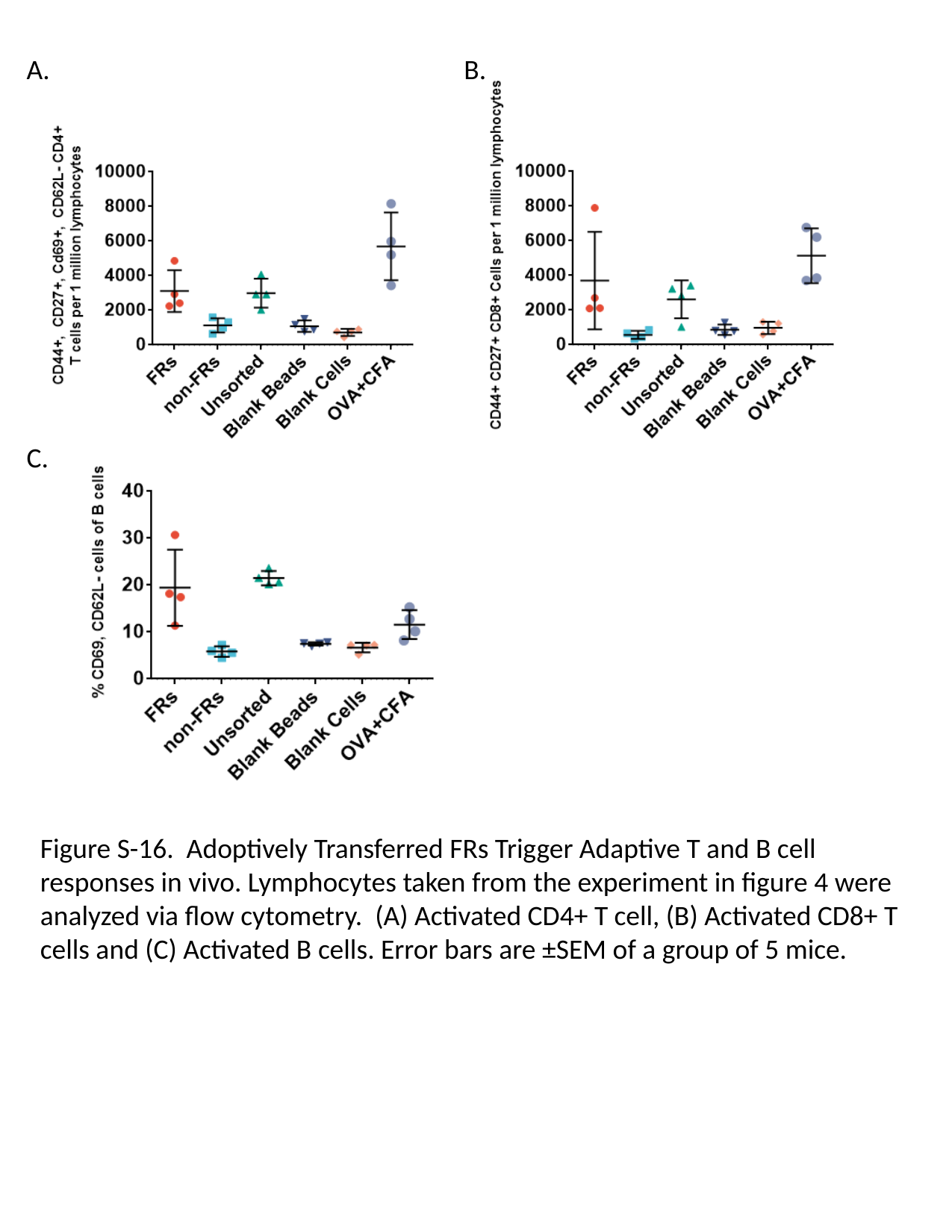

A.
B.
C.
Figure S-16. Adoptively Transferred FRs Trigger Adaptive T and B cell responses in vivo. Lymphocytes taken from the experiment in figure 4 were analyzed via flow cytometry. (A) Activated CD4+ T cell, (B) Activated CD8+ T cells and (C) Activated B cells. Error bars are ±SEM of a group of 5 mice.

### Slide 18
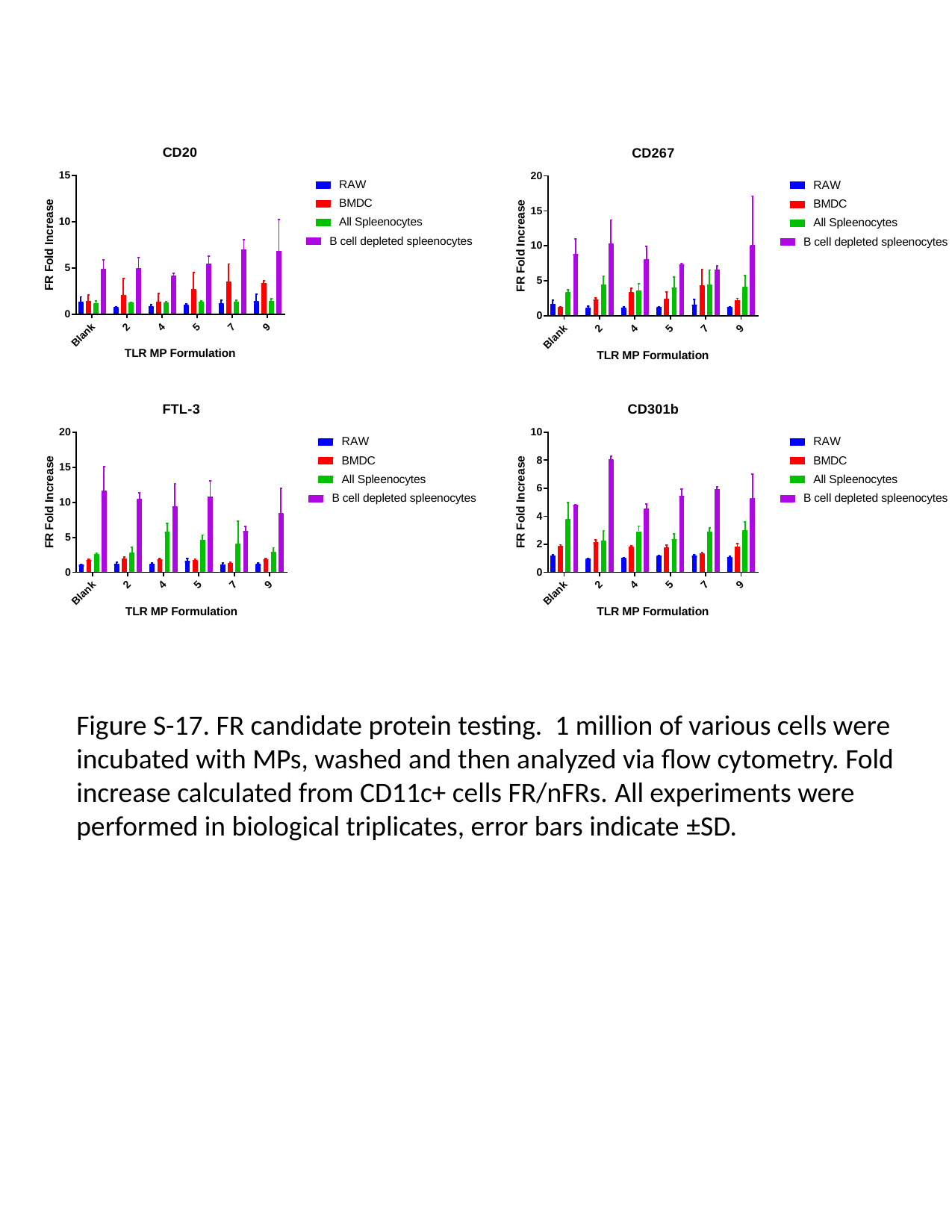

Figure S-17. FR candidate protein testing. 1 million of various cells were incubated with MPs, washed and then analyzed via flow cytometry. Fold increase calculated from CD11c+ cells FR/nFRs. All experiments were performed in biological triplicates, error bars indicate ±SD.

### Slide 19
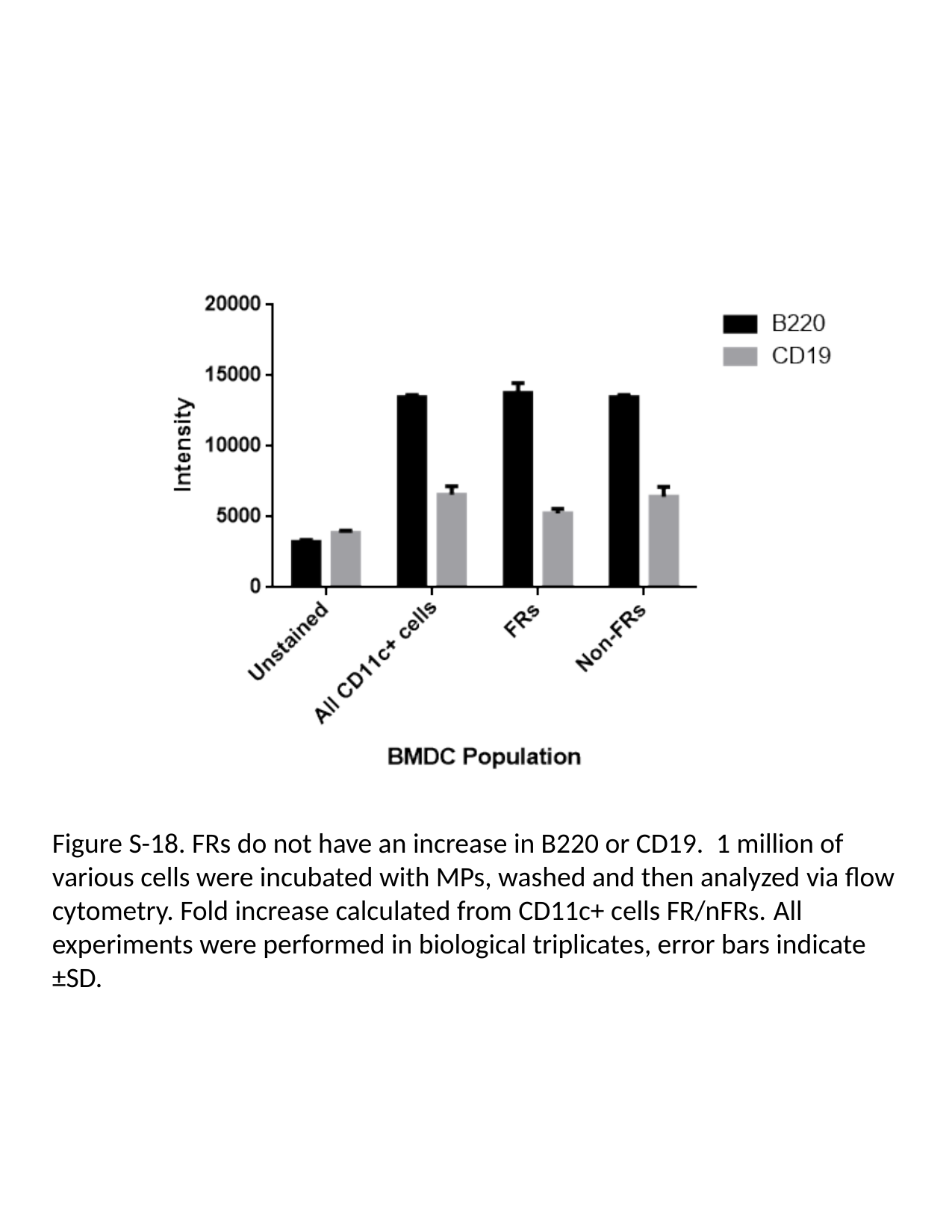

Figure S-18. FRs do not have an increase in B220 or CD19. 1 million of various cells were incubated with MPs, washed and then analyzed via flow cytometry. Fold increase calculated from CD11c+ cells FR/nFRs. All experiments were performed in biological triplicates, error bars indicate ±SD.

### Slide 20
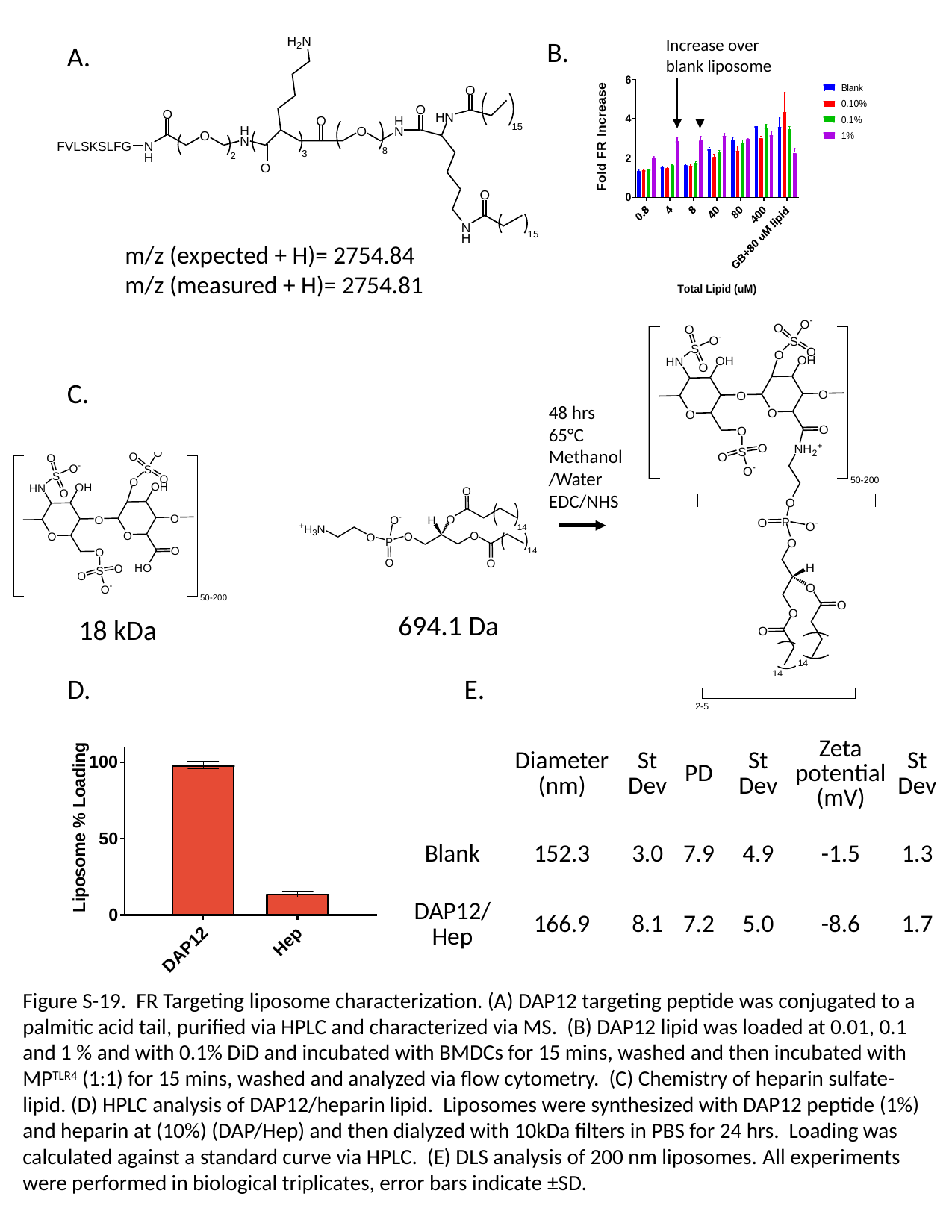

Increase over blank liposome
B.
A.
m/z (expected + H)= 2754.84
m/z (measured + H)= 2754.81
C.
48 hrs
65°C
Methanol/Water
EDC/NHS
694.1 Da
18 kDa
D.
E.
| | Diameter (nm) | St Dev | PD | St Dev | Zeta potential (mV) | St Dev |
| --- | --- | --- | --- | --- | --- | --- |
| Blank | 152.3 | 3.0 | 7.9 | 4.9 | -1.5 | 1.3 |
| DAP12/Hep | 166.9 | 8.1 | 7.2 | 5.0 | -8.6 | 1.7 |
Figure S-19. FR Targeting liposome characterization. (A) DAP12 targeting peptide was conjugated to a palmitic acid tail, purified via HPLC and characterized via MS. (B) DAP12 lipid was loaded at 0.01, 0.1 and 1 % and with 0.1% DiD and incubated with BMDCs for 15 mins, washed and then incubated with MP^TLR4^ (1:1) for 15 mins, washed and analyzed via flow cytometry. (C) Chemistry of heparin sulfate-lipid. (D) HPLC analysis of DAP12/heparin lipid. Liposomes were synthesized with DAP12 peptide (1%) and heparin at (10%) (DAP/Hep) and then dialyzed with 10kDa filters in PBS for 24 hrs. Loading was calculated against a standard curve via HPLC. (E) DLS analysis of 200 nm liposomes. All experiments were performed in biological triplicates, error bars indicate ±SD.

### Slide 21
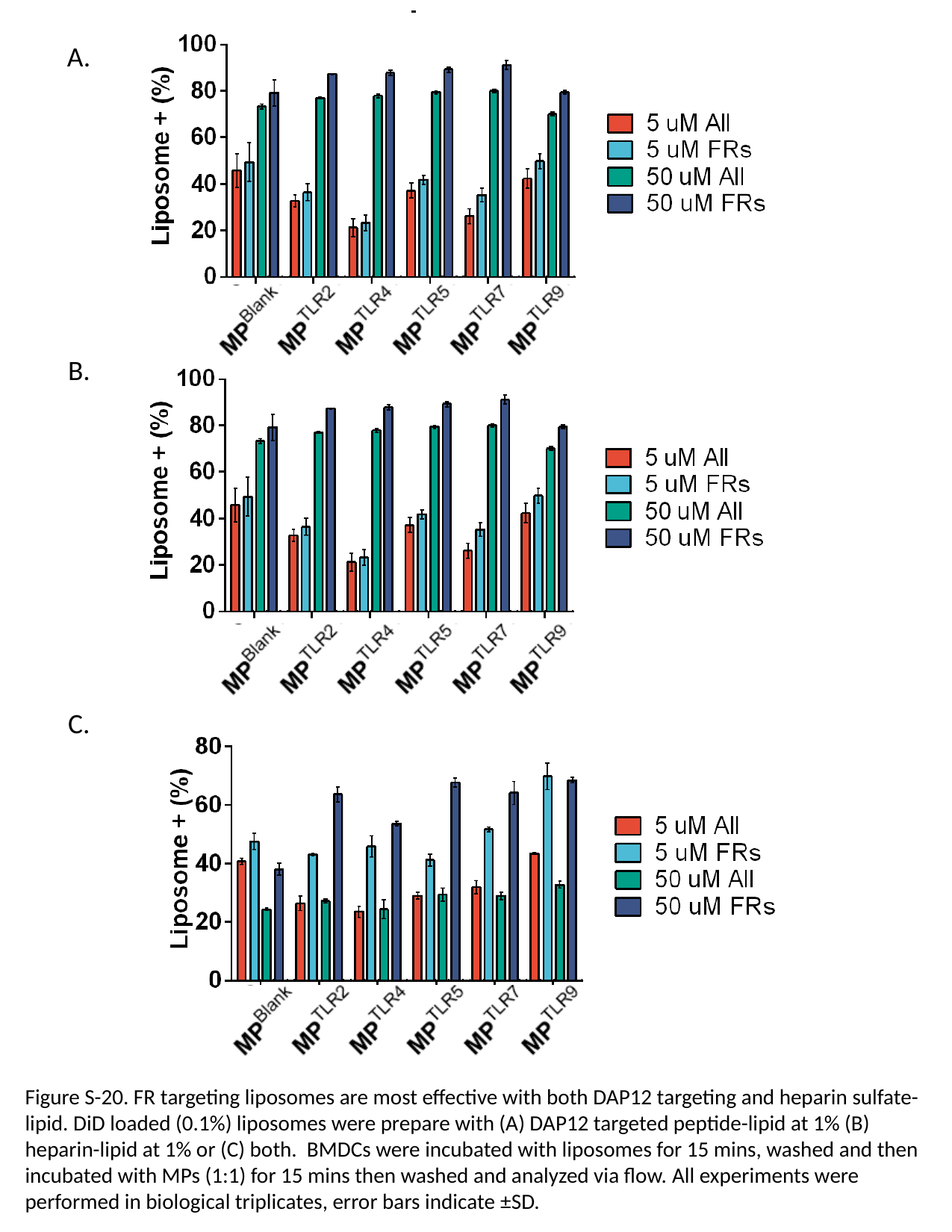

A.
B.
C.
Figure S-20. FR targeting liposomes are most effective with both DAP12 targeting and heparin sulfate-lipid. DiD loaded (0.1%) liposomes were prepare with (A) DAP12 targeted peptide-lipid at 1% (B) heparin-lipid at 1% or (C) both. BMDCs were incubated with liposomes for 15 mins, washed and then incubated with MPs (1:1) for 15 mins then washed and analyzed via flow. All experiments were performed in biological triplicates, error bars indicate ±SD.

### Slide 22
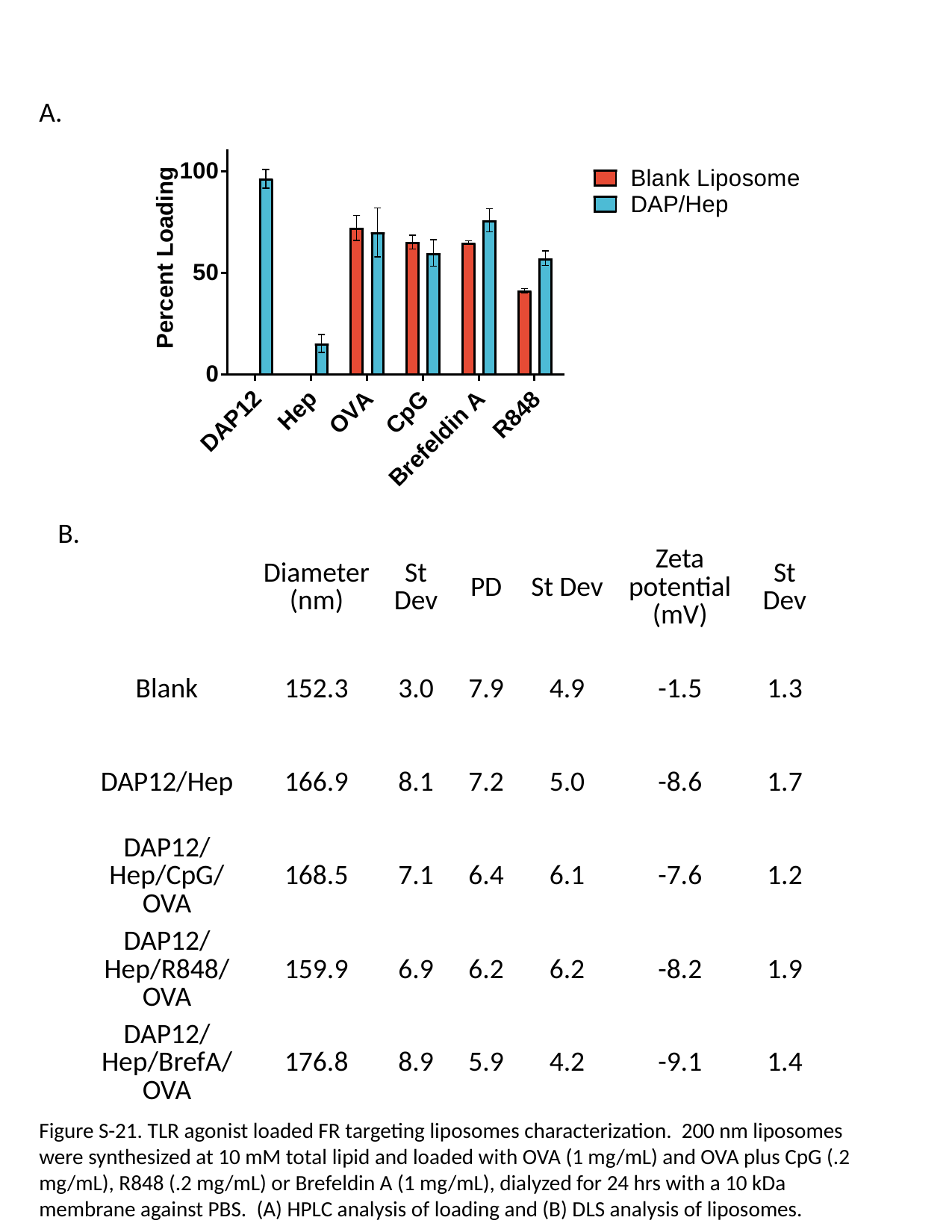

A.
B.
| | Diameter (nm) | St Dev | PD | St Dev | Zeta potential (mV) | St Dev |
| --- | --- | --- | --- | --- | --- | --- |
| Blank | 152.3 | 3.0 | 7.9 | 4.9 | -1.5 | 1.3 |
| DAP12/Hep | 166.9 | 8.1 | 7.2 | 5.0 | -8.6 | 1.7 |
| DAP12/Hep/CpG/OVA | 168.5 | 7.1 | 6.4 | 6.1 | -7.6 | 1.2 |
| DAP12/Hep/R848/OVA | 159.9 | 6.9 | 6.2 | 6.2 | -8.2 | 1.9 |
| DAP12/Hep/BrefA/OVA | 176.8 | 8.9 | 5.9 | 4.2 | -9.1 | 1.4 |
Figure S-21. TLR agonist loaded FR targeting liposomes characterization. 200 nm liposomes were synthesized at 10 mM total lipid and loaded with OVA (1 mg/mL) and OVA plus CpG (.2 mg/mL), R848 (.2 mg/mL) or Brefeldin A (1 mg/mL), dialyzed for 24 hrs with a 10 kDa membrane against PBS. (A) HPLC analysis of loading and (B) DLS analysis of liposomes.

### Slide 23
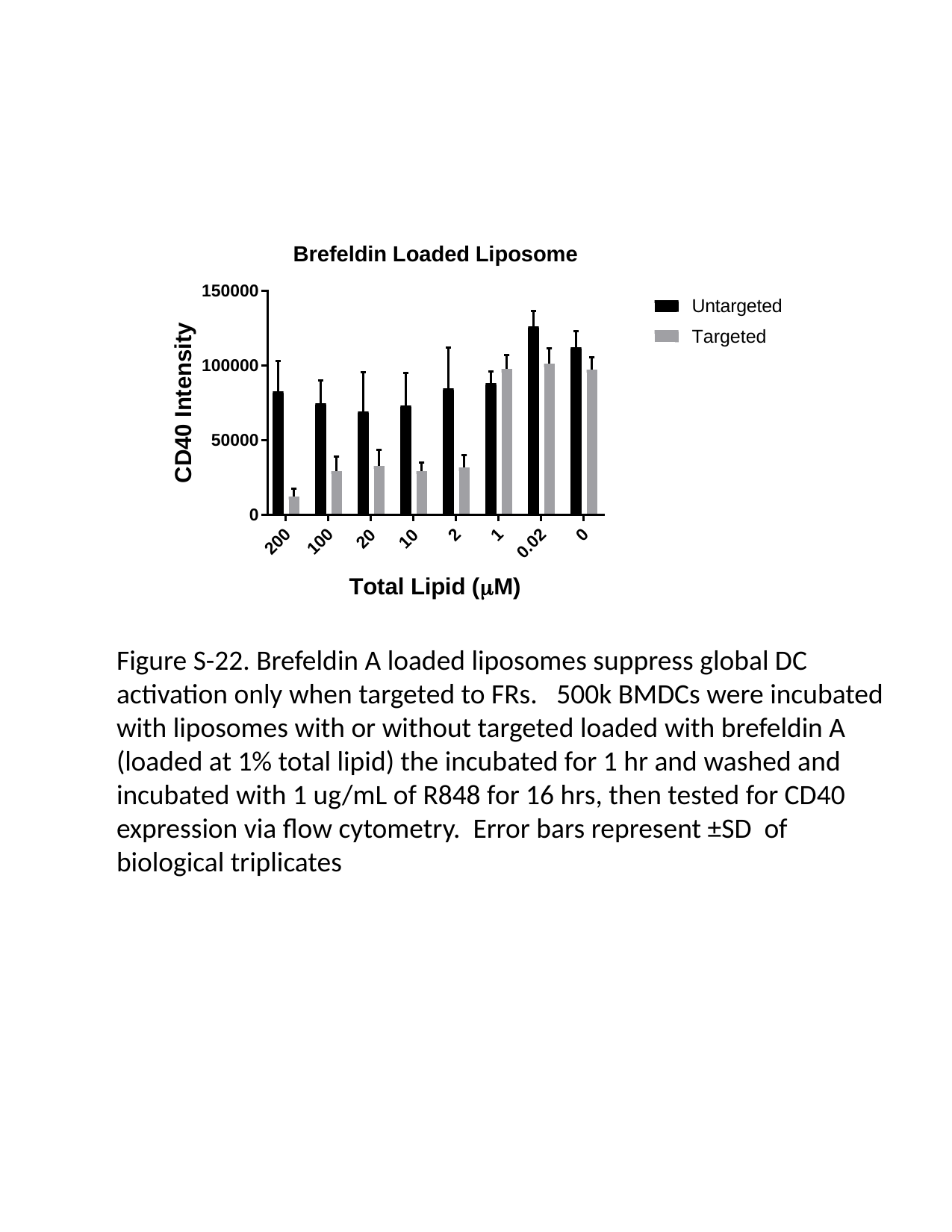

Figure S-22. Brefeldin A loaded liposomes suppress global DC activation only when targeted to FRs. 500k BMDCs were incubated with liposomes with or without targeted loaded with brefeldin A (loaded at 1% total lipid) the incubated for 1 hr and washed and incubated with 1 ug/mL of R848 for 16 hrs, then tested for CD40 expression via flow cytometry. Error bars represent ±SD of biological triplicates

### Slide 24
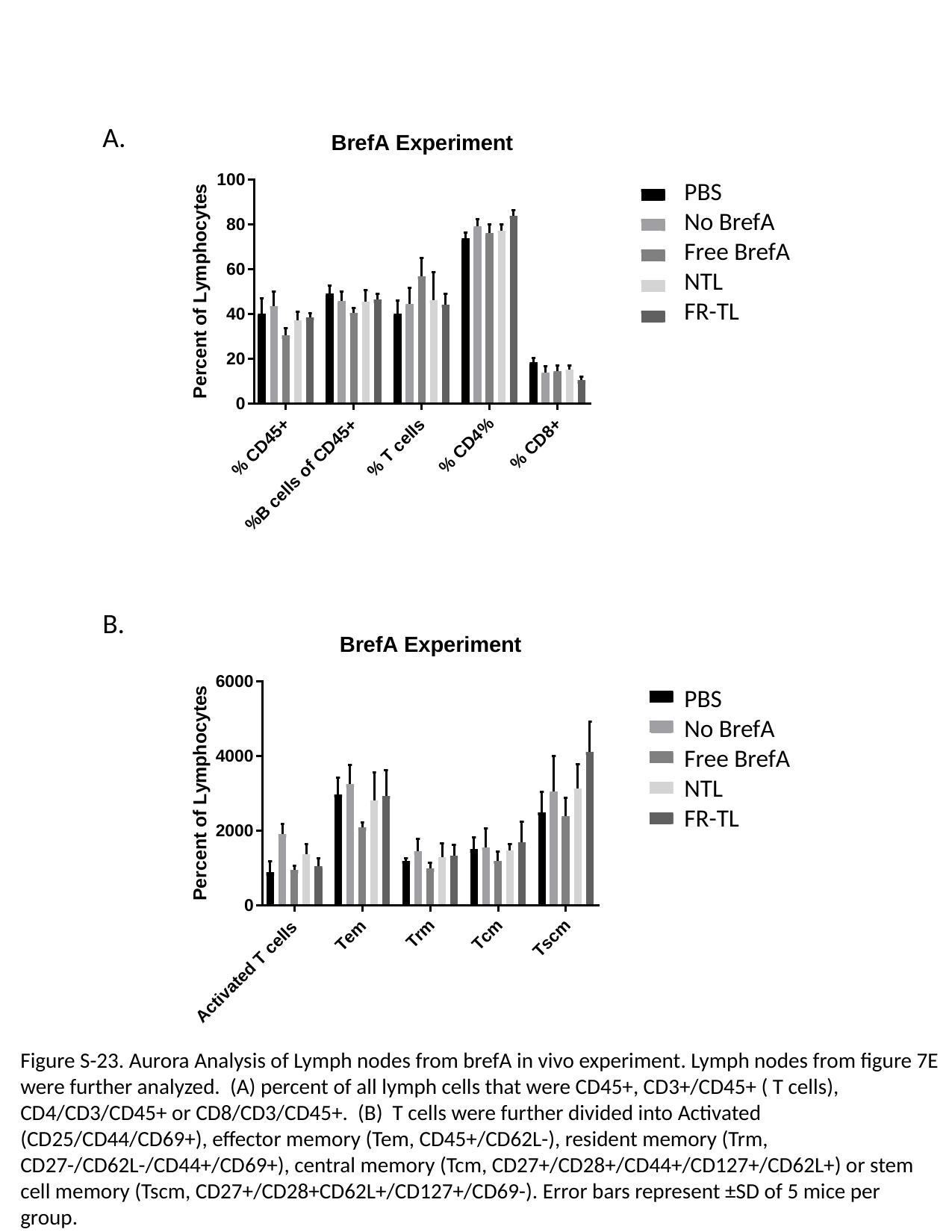

A.
PBS
No BrefA
Free BrefA
NTL
FR-TL
B.
PBS
No BrefA
Free BrefA
NTL
FR-TL
Figure S-23. Aurora Analysis of Lymph nodes from brefA in vivo experiment. Lymph nodes from figure 7E were further analyzed. (A) percent of all lymph cells that were CD45+, CD3+/CD45+ ( T cells), CD4/CD3/CD45+ or CD8/CD3/CD45+. (B) T cells were further divided into Activated (CD25/CD44/CD69+), effector memory (Tem, CD45+/CD62L-), resident memory (Trm, CD27-/CD62L-/CD44+/CD69+), central memory (Tcm, CD27+/CD28+/CD44+/CD127+/CD62L+) or stem cell memory (Tscm, CD27+/CD28+CD62L+/CD127+/CD69-). Error bars represent ±SD of 5 mice per group.

### Slide 25
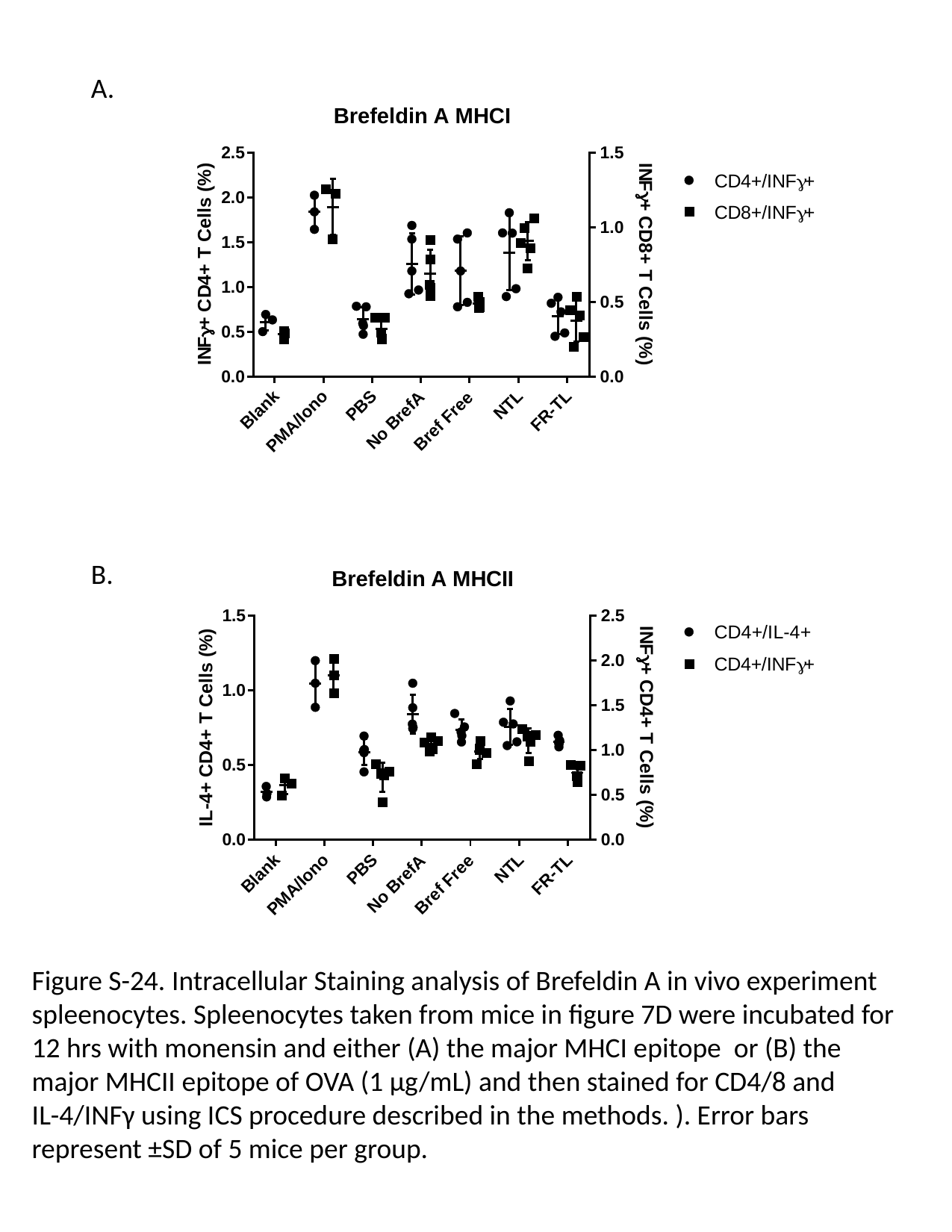

A.
B.
Figure S-24. Intracellular Staining analysis of Brefeldin A in vivo experiment spleenocytes. Spleenocytes taken from mice in figure 7D were incubated for 12 hrs with monensin and either (A) the major MHCI epitope or (B) the major MHCII epitope of OVA (1 µg/mL) and then stained for CD4/8 and IL-4/INFγ using ICS procedure described in the methods. ). Error bars represent ±SD of 5 mice per group.

### Slide 26
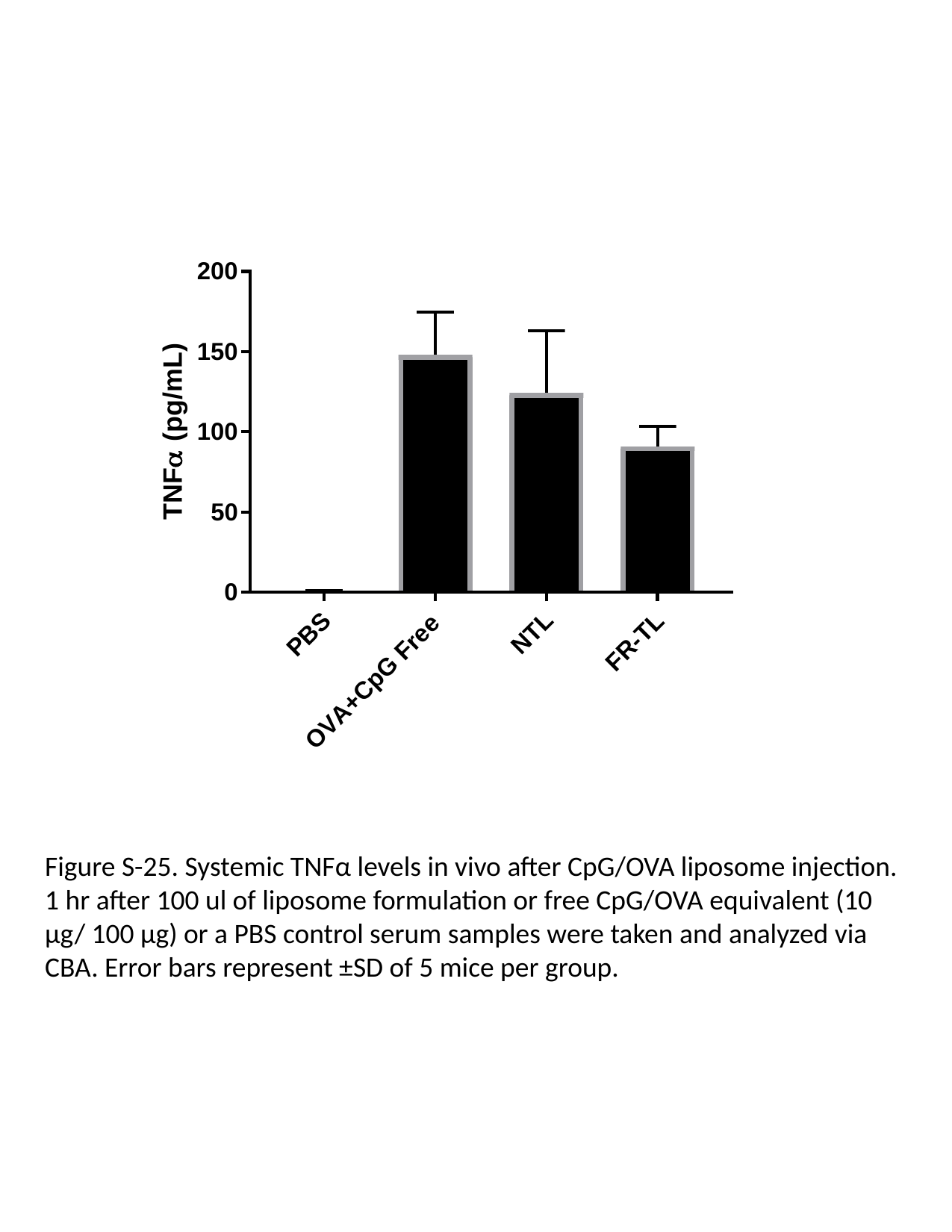

Figure S-25. Systemic TNFα levels in vivo after CpG/OVA liposome injection. 1 hr after 100 ul of liposome formulation or free CpG/OVA equivalent (10 µg/ 100 µg) or a PBS control serum samples were taken and analyzed via CBA. Error bars represent ±SD of 5 mice per group.

### Slide 27
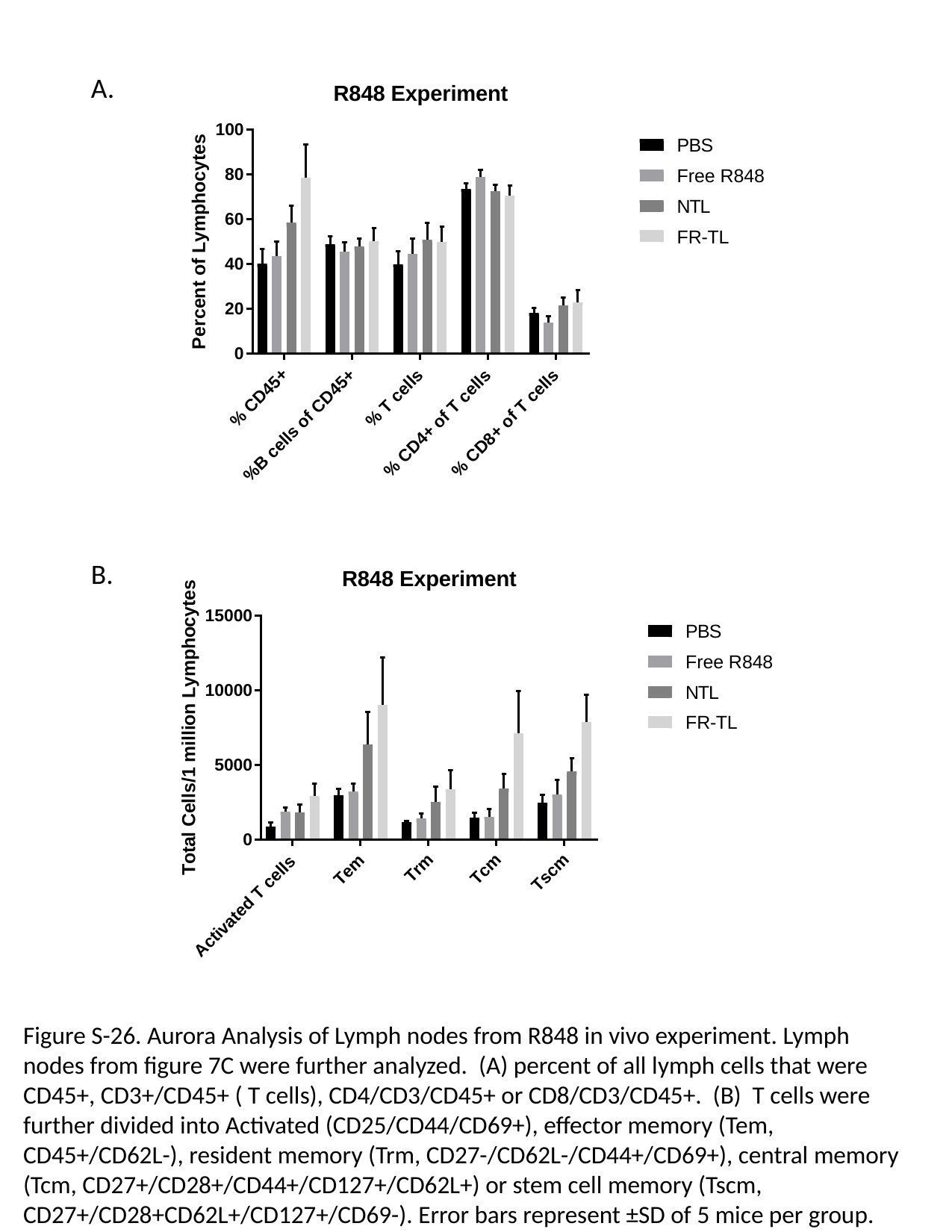

A.
B.
Figure S-26. Aurora Analysis of Lymph nodes from R848 in vivo experiment. Lymph nodes from figure 7C were further analyzed. (A) percent of all lymph cells that were CD45+, CD3+/CD45+ ( T cells), CD4/CD3/CD45+ or CD8/CD3/CD45+. (B) T cells were further divided into Activated (CD25/CD44/CD69+), effector memory (Tem, CD45+/CD62L-), resident memory (Trm, CD27-/CD62L-/CD44+/CD69+), central memory (Tcm, CD27+/CD28+/CD44+/CD127+/CD62L+) or stem cell memory (Tscm, CD27+/CD28+CD62L+/CD127+/CD69-). Error bars represent ±SD of 5 mice per group.

### Slide 28
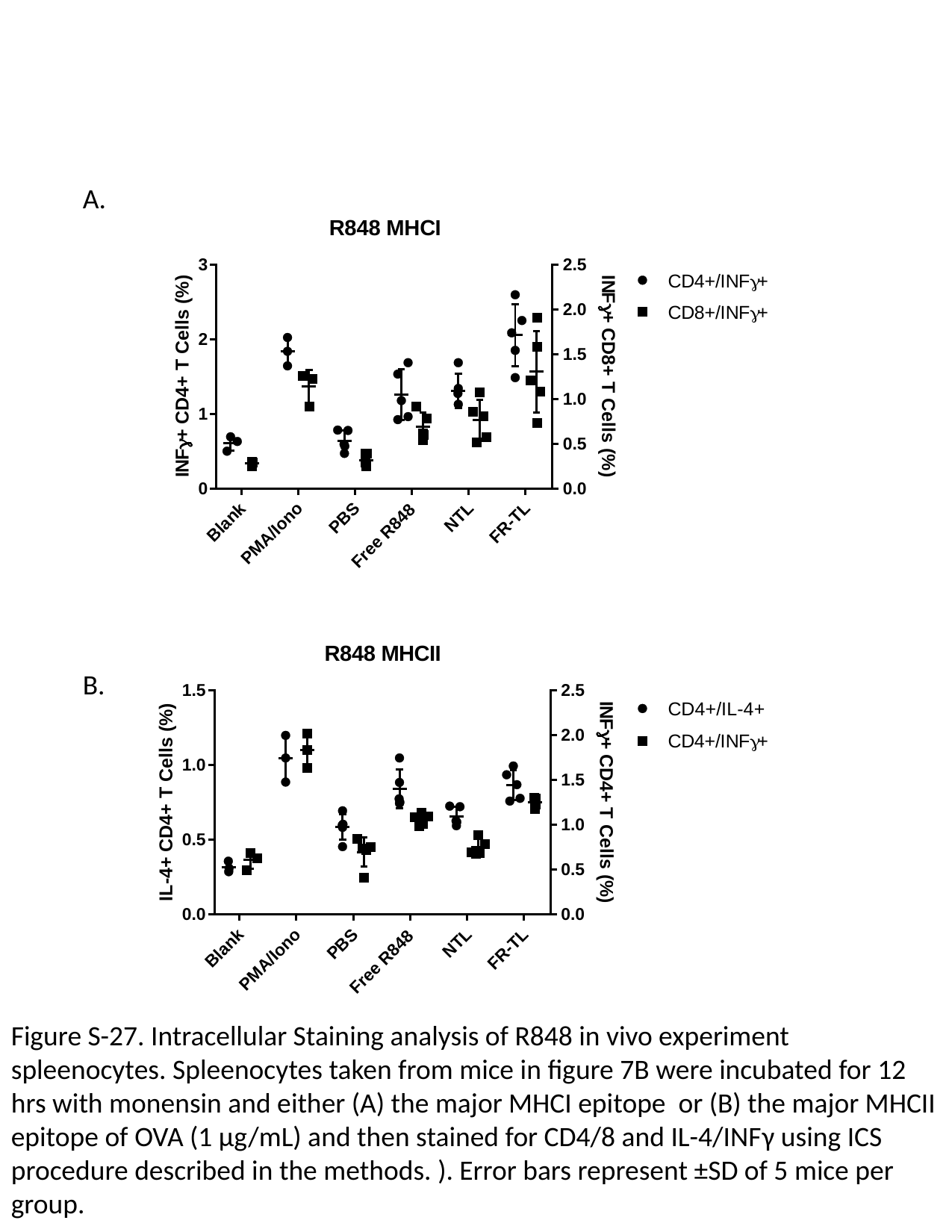

A.
B.
Figure S-27. Intracellular Staining analysis of R848 in vivo experiment spleenocytes. Spleenocytes taken from mice in figure 7B were incubated for 12 hrs with monensin and either (A) the major MHCI epitope or (B) the major MHCII epitope of OVA (1 µg/mL) and then stained for CD4/8 and IL-4/INFγ using ICS procedure described in the methods. ). Error bars represent ±SD of 5 mice per group.

### Slide 29
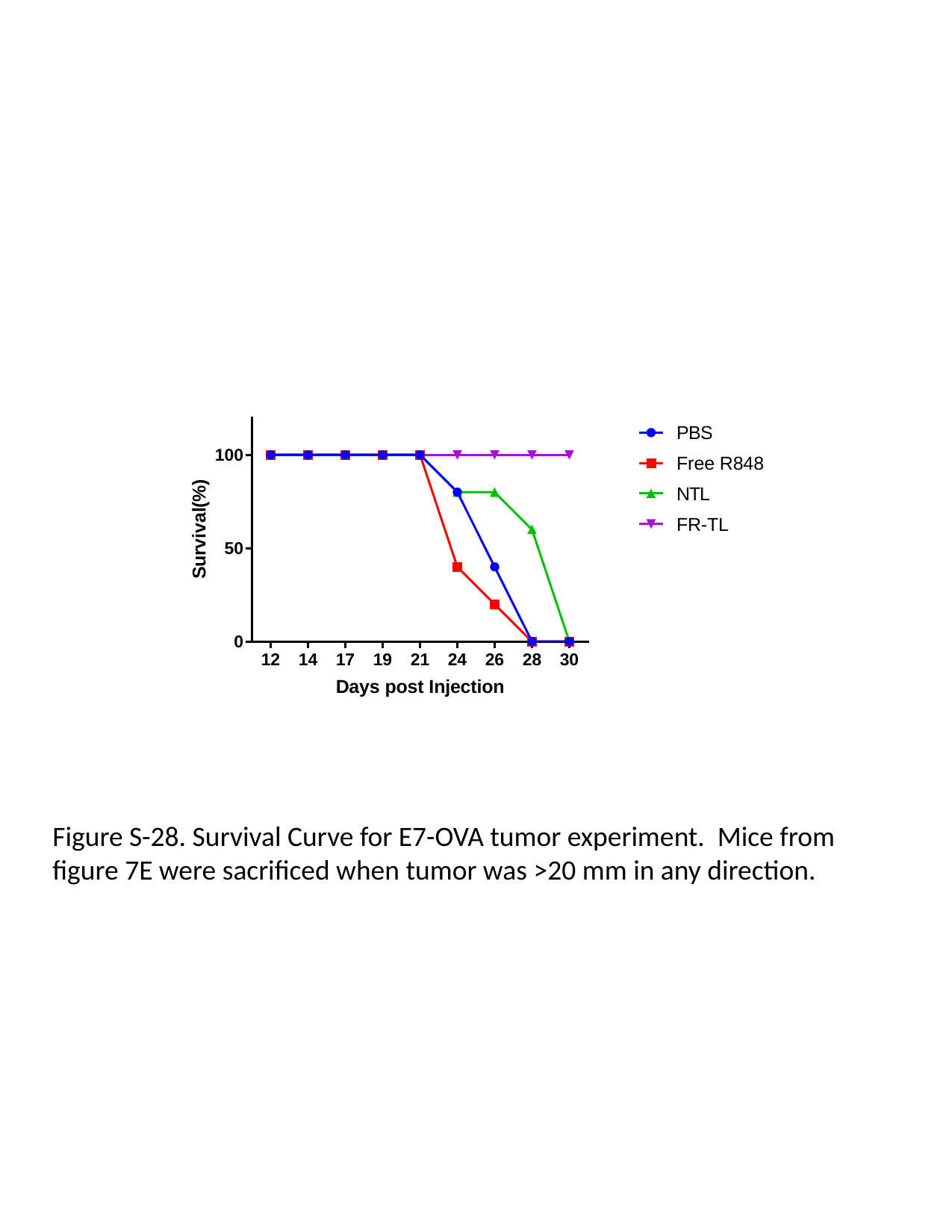

Figure S-28. Survival Curve for E7-OVA tumor experiment. Mice from figure 7E were sacrificed when tumor was >20 mm in any direction.
