## Supplemental Materials and Methods for "Identification, Characterization, and Targeting of a Rare and Temporal Dendritic Cell State that Facilitates Adaptive Immune Responses"

***Materials***

All chemicals were purchased from Sigma unless noted. Lipopolysaccaride from E. coli (LPS), Flagellin from Bacillus subtilis (FLA), Lipoteichoic acid from S. aureus (LTA), R848, CpG 1826, Ovalbumin (OVA) were purchased from Invivogen. The TLR7 agonist 2BXY was synthesized in house according to published protocols.^1^ Thiol functionalized CpG 1826 (GpG-SH) was purchased as a custom order from Integrated DNA technologies. BCA assay kit was purchased from Thermo. Cytoperm/CytoFix Kit plus GolgiPlug was purchased from BD Biosciences. All antibodies were purchased from BD Biosciences. All peptide synthesis reagents were purchased from EMD Millipore. Quanti Blue assay solution was purchased from Invivogen. Anti-OVA IgG ELISA kit was purchased from Alpha Diagnostic International.

***Methods***

*Silica-silane coated Polystyrene Microparticle (MP) synthesis:* PS MPs were synthesized and coated with a silica coating using a procedure from Moser et al.^2^ Briefly, uniform, spherical, 2 μm diameter polystyrene microparticles were synthesized via controlled styrene polymerization. 2 g of polyvinylpyrrolidone, MW 40,000 and styrene (20 g), washed with NaOH and dried with MgSO4, was dissolved in EtOH (250 mL) and purged with nitrogen. AIBN (0.2 g) was added, the mixture stirred at 70 °C and 200 rpm for 24 hrs. Mixture was purified by centrifugation (5000 RPM for 5 minutes, followed by washing 3X in 30 mL of EtOH to remove residual monomer, initiator, and stabilizer). The surfaces of MPs were modified with reactive thiol groups via Pickering emulsion reaction. A mixture of Cyclohexane (45 mL), n-hexanol (10.8 mL), endotoxin free water (2 mL) and Triton X-114 (10.8 mL) were placed in a round-bottom flask and sonicated for 20 min. Particles (0.2 g) were added and the suspension was sonicated for 40 min. TEOS (400 μL) was added dropwise followed by 14 M aqueous ammonia (1.2 mL). The resulting solution was stirred for 30 min RT. Subsequently, 3-Mercaptosilane (200 μL) was added dropwise and stirred for 6 h. TEOS-mercaptosilane copolymer coated particles were then pelleted at 3400 rpm for 30 min and washed 3× with EtOH. Particles were dried at 70 °C and stored at 4 °C.

*TLR agonist Surface Functionalization:* Thiol bearing MPs were functionalized with MPLA and Pam_2_ using thiol-maleimide chemistry. First MPs (5 mg) were swelled in ACN (500 uL) for 30 mins under sonication, then 1 mg of FITC in 500 uL of ACN was added for a final concentration of 1 mg/mL for 30 mins. MPs were then centrifuged for 1 min (5000 rcf), supernatant removed and washed 3x with PBS. For MPs conjugated with agonists to TLR2, 5, 7 and 9: FITC labeled MPs were dissolved in 500 uL of PBS, then 5 mg of Bismaleimide-PEG_3_ was added and allowed to sonicate for 30 minutes, then washed 3x in PBS. During this LTA, FLA, 2BXY or CpG-SH (0.1 mg/mL) was incubated for 5 mins with Trauts reagent (1 mg/mL) in 500 uL of PBS. After washing, maleimide bearing MPs were incubated with thiol functionalized with either Trauts + LTA, FLA, 2BXY or CpG-SH (500 uL) and sonicated for 30 mins, then washed 10 times, 3x with PBS, 4x with PBS with 0.1 tween 20, and 3x with PBS. For TLR9 MPs, 1 mg of LPS was dissolved in 400 uL of DMF and 10 ug of p maleimidophenyl isocyanate added and allowed to stir overnight under argon. Then PBS (400 uL) was added followed by 2,2’-(ethylenedioxy)diethanethiol (4 mg) and allowed to stir for 12 hrs. 100 uL of this mixture was added to the MP with 400 uL of PBS for 30 mins and similarly washed. After conjugation, all MPs were diluted in PBS + 0.05 %wt tween20 at 20, 50 and 100X and then number of MPs in 10 uL counted using flow cytometry to determine a concentration. MP solutions were diluted to a concentration of 1 million MPs per uL of PBS and stored at 4ºC until use.

*Particle Concentration Determination via Flow Cytometry-* The concentration of particle in a given volume was determined using flow cytometry (NovoCyte flow cytometer, ACEA Biosciences, Inc). After the final wash, MPs were rehydrated in 500 μL of PBS, then diluted down in PBS +0.01 %wt tween 20 by 50, 100 and 200 fold. 10 μL of these dilutions were analyzed via flow cytometry in triplicate, gating on FITC signal to remove noise. The MP concentration in the stock was then calculated using linear regression from the three dilutions. MP stocks were then diluted down with PBS to a final concentration of 1 million MPs/μL.

*Synthesis and Purification of DAP12 targeting peptide lipid* The peptide-lipid for liposome encapsulation was adapted from a paper by Stefanick et al.^3^ Synthesis was performed using a Liberty Blue™ automated peptide synthesizer using the schematic in Figure S-17. Rink amide resin (100-200 mesh, 0.55 mmole/g, 0.05 mg) was weighed out into a solid-phase peptide synthesizer reaction vessel. The peptide (FVLSKSLFG) was constructed with standard Fmoc protected peptides (0.2M in DMF) from the C terminus to the N terminus. Then a Fmoc-EG2-COOH was conjugated followed by three lysines, EG8 linker, a Fmoc-Lys(Fmoc)-OH and palmitic acid. Deprotection was performed using 20% piperidine in DMF. Coupling was performed after activation with diisopropylcarbodiimide (DIC) (0.5M in DMF) in the presence of Ethyl cyanohydroxyiminoacetate (oxyma) (1M in DMF). All other couplings were done at 90 ^o^C for 5 min. All reactions and subsequent washes were performed in DMF. After the synthesis was completed, the resin was transferred into a Bio-Rad Poly-Prep chromatography column.

Global deprotection was achieved by agitating the resin in trifluoroacetic acid (TFA)/3,6-dioxa-1,8-octanedithiol (DODT)/triisopropylsilane (TIPS)/H2O (8.5:0.5:0.5:0.5) for 2 h. The peptide was precipitated by adding the cleavage cocktail filtrate to 30 mL diethyl ether in a 50 mL centrifuge tube pre-cooled to -78 ^o^C. The precipitate was collected by centrifuge (4000 XG for 5 min). The precipitate was dissolved in 50% CH_3_CN in 0.1%TFA) and filtered through a 0.45 µm syringe filter. Purification was performed using reversed-phase HPLC C8 column (gradient elution with 30–90% CH_3_CN /0.1% TFA over 20min). Pure fractions were pooled together and the peptide was recovered through lyophilization and redissolved in MeOH. LC-MS (m/z) 2754.84 [M+H] +

*Synthesis and Purification of Heparin Sulfate Lipid* This procedure was adapted from a paper by Kim et al.^4^ 300 mg Heparin Sulfate (18 kDa) was dissolved in water (10 mL) and mixed with DPPE (60 mg) dissolved in IPA (10 mL). 138 mg EDC and 69 mg of NHS were added. The mixture was stirred constantly at 65°C for 20 hrs. After reaction, the mixture was rotoevaporated to remove excess IPA, lyophilized, redissolved in 5 mL of water, filtered and dialyzed against water in a 3500 Da cutoff dialysis cassette for 48 hrs. Then the purified Heparin-lipid was analyzed via LC-MS, confirming between 2-5 conjugation of DPPE per heparin chain, lyophilized again and redissolved at 10 mg/mL in MeOH.

*Liposome Synthesis* Liposomes were synthesized via membrane extrusion method using a setup from Avanti polar lipids and 200 nm extrusion filters. DSPC, PEG2000 PE, Cholesterol, and/or Did, Heparin-lipid, DAP12 peptide lipid and any TLR agonists/ Brefeldin A were combined in 1 mL methanol added, dried via lyophilization, and rehydrated in PBS to make a 10 mM total lipid, 200 uL solutions. OVA was added during rehydration. Solutions were gently rotated at 67°C and then passed through a 70°C 200 nm filter 5 times. The liposome solution was dialyzed against PBS with a 3500 Da filter for 24 hrs.

*LC-MS Analysis of Liposome Loading* Liposomes loaded with DiD, DAP12 peptide, Heparin, brefeldin A, CpG, R848, OVA or any combinations of these were tested for loading via LC-MS analysis. Stock solutions of liposomes were diluted down to 1 mM total lipid and then 5 uL of this solution was injected on a C8 analytical HPLC column with a gradient of 10-90% ACN in 20 mins and the effluent observed by an Agilent 6135BAR LCMS XT mass spectrometer and a diode array detector at 220 nm. Signal from various compounds in liposome formulations were compared to standard curves at 220 nm.

*Dynamic Light Scattering Analysis of Liposomes* Hydrodynamic diameter of liposomes was confirmed using a Wyatt Möbiuζ DLS/ELS at 100 uM total liposome concentration.

*SEM of Functionalized Particles*: Scanning electron microscopy (SEM) and energy-dispersive spectroscopy (EDS) of the particles was performed using an FEI Quanta 3D FEG dual beam (SEM/FIB) equipped with Inca EDS (Oxford Instruments). High-resolution images were taken with an FEI Magellan 400 XHR SEM particle samples were dried under vacuum for 24 h, mounted on carbon tape, and sputter coated (South Bay Technologies) with approximately 2–4 nm of Au/Pd 60:40 or Ir.

*Cell Culture* RAW 264.7 Cells were cultured at 1 million cells per mL in DMEM+ Penicillin/Streptomycin+ 10% HIFBS and split every 2-3 days.

*BMDC Cell Culture* BMDCs were cultured according to a previously published protocol. Cells were used between 5-7 days after isolation.^5^

*Animals* 6 week C57BL/6 female mice were purchased from Jackson Laboratory and housed and treated according to our approved IACUC protocol.

*Spleeyocyte/Lymph Node/Footpad extraction* Spleeyocyte/Lymph Node/Footpad were dissected into 1 mm sized portions and placed in disassociation media (0.5 mg/mL collagenase D, 0.1 mg/mL DNAase I in RPMI) for 30 mins at room temp, then incubated at 37°C for 30 mins, then passed through a 70 um filter. Footpads were incubated for 2 hrs rather than 30 mins. Spleenocytes were treated with RBC lysis buffer (Invitrogen) prior to final wash.

*RAW Blue NF-κB Assay:* RAW-Blue NF-κB cells (Invivogen) were passaged and plated in a 96 well plate at 100k cells/well in 180 μL DMEM containing 10% HIFBS. Cells were incubated at 37 °C and 5% CO_2_ for 24 h. 100 ul of cells were incubated with varying ratios of MPs at 37 °C and 5% CO2 for 18 h. After 18 h, 20 μL of the cell supernatant was placed in 180 μL freshly prepared QuantiBlue (Invivogen) solution and incubated at 37 °C/5% CO2 for up to 2 h. The plate was analyzed every hour using a Multiskan FC plate reader (Thermo Scientific) and absorbance was measured at 620 nm *BCA Assay-* This was performed according to manufacturer’s instruction (Thermo Fischer) with some modifications. 100 million beads were incubated with BCA solution and reacted for 30 mins at 60 ºC then analyzed every hour using a Multiskan FC plate reader (Thermo Scientific) and absorbance was measured at 562 nm and compared to a standard curve of modified MPLA or Pam_2_ after subtracting a background of maleimide modified MP.

*Flow Cytometry* Most flow cytometry in this study was performed on a ACEA NovaCyte Flow cytometer (6 channels, 2 laser). 1 million cells per sample were treated with liposomes/MPs/antibodies, washed and placed in HBSS+ 2% HIFBS +0.1 mM EDTA. Samples were gated on FCS and SCC for live and single cells. Samples were compensated based on sample with single stains and calculated using FlowJo.

*Aurora Spectral Flow Analysis* A 5 laser Aurora spectral flow cytometer was used for phenotyping DCs and T cells. Single stained compensation controls were run first and then unmixed using the Aurora spectral flow software. Data was further analyzed in FlowJo.

*Image Stream MP Uptake Analysis for TNFα Expression:* 1 million BMDCs, RAWs or Spleenocytess were incubated with 1 million MPs in 1 mL of cell culture media (DMEM supplanted with 10% HIFBS) for 1 hour. After 1 hr, protein export was inhibited using a GolgiPlug Kit (BD Biosciences) with Brefeldin A according to manufacturer’s instructions for 16 hrs at 37 ºC under 5% CO_2_. Cells were then washed and fixed and permeabilized with a BD Cytofix/Cytoperm Plus Kit (BD Biosciences). Cells were stained with a solution of anti-TNFα (1:500 dilution) and their nuclei stained using Hoechst 33342 Solution (2 μM final concentration) in permeabilization buffer for 1 hr. Cells were washed 3x in PBS with 2% HIFBS, concentrated into a 20 μL volume and analyzed with ImageStream Flow Cytometry. Each MP condition was performed in triplicate, analyzing >100,000 cells per run. MPs were identified using the “Particle Count” wizard in the IDEAS software and compared to TNFα intensity per cell.

*Image Stream MP Uptake Analysis for NF-κB Expression:* 1 million BMDCs, RAWs or Spleenocytes were incubated with 1 million MPs in 1 mL of cell culture media (DMEM supplanted with 10% HIFBS) for 15 minutes. Immediately cells were fixed using ice cold Cytofixation Buffer (BD Biosciences) for 15 minutes. Cells were then fixed in PBS +0.04% triton X for 3 minutes, immediately spun down (400 RCF, 5 minutes), supernatant removed and washed with CytoPerm Buffer (BD Biosciences) 2 times (200 μL per wash). Cells were then stained with Rabbit anti-NF-κB p65 (1:500 dilution) for 1 hr on ice, washed 3x, then stained with a secondary goat anti-rabbit AF647 (1:1000 dilution) for 1 hr on ice, washed, concentrated into a 20 μL volume of PBS with 2% HIFBS and analyzed with ImageStream Flow Cytometry. Each MP condition was performed in triplicate, analyzing >100,000 cells per run. MPs were identified using the “Particle Count” wizard in the IDEAS software and compared to NF-κB nuclear colocalization using the “Colocalization” wizard in the IDEAS software.

*Imagestream Data Analysis:* ImageStream data was first analyzed in the IDEAS software (Amnis) for nuclear colocalization and particle counting using built-in analysis wizards. Single Cell data was then exported into Graphpad Prism 6 software for further analysis. Cell data was divided into the following categories: 0 MP, 1 MP, 2 MP, 3 MP, 4 MP or 5 MP or >5 MPs.

*Microscopy* BMDCs, Spleenocytes or RAWs were analyzed with a SP5 two photon confocal microscope. 100k cells were allowed to attach to the bottom of a 96 well plate in their respective cell culture media. The next day, cells were washed with HBSS then either incubated with DiD containing liposomes for 15 mins, washed and incubated with MPs for 15 mins ord incubated with MPs for 15 mins, washed and then incubated with antibodies. Cells were washed and placed in fluorobrite media (Gibco) with 10% HIFBS +1:2000 dilution of Hoest then analyzed by microscopy using relevant wavelengths/filters. Single images were taken using a 60x lens.

*Cell Sorting* BMDCs, Spleenocytes or RAWs were sorted on an Aria Fusion 5-18, AriaIIIu 4-15 or AriaII 4-15 cell sorter. For sorting, cells were sometimes treated with or without Brefeldin A, then incubated with MPs for 15 mins, washed, scrapped, washed again and diluted to 20 million cells per mL in RPMI. Cells were gated on live and single cells by FSC and SCC and sorted into FR (top 5% of FITC signal from MPs) and nFR (bottom 90% of FITC signal from MPs). Cells were then immediately placed on ice, spun down at 4°C and placed in cell culture media for subsequent experiments. For kinetic experiments, unsorted cell controls were also run through sorter but only gated on live and single cells.

*Cytokine Bead Array* a Mouse Inflammation CBA kit was purchased from BD Biosciences and used according to the manufacturer’s instructions. Mouse blood was spun down at 10000 g for 10 mins to remove cells and the supernatant tested undiluted. Supernatant from cell culture experiments was also used with no dilution.

*First Responder Cytokine Analysis* BMDCs were incubated at a 1:1 ratio with MP^TLR-4^ for 15 minutes and the FRs and nFRs were isolated via FACS. The cells were washed and resuspended at 1 million cells/mL in culture media (10% HIFBS in RPMI). The cells were incubated at 37°C and 5% CO2 for 1 h. The supernatant was collected and stored at -80°C until the cytokines were profiled using a mouse inflammation CBA kit (BD Biosciences).

*First Responder Kinetic Analysis* Sorted BMDC FRs or nFRs cells (BMDCs incubated 1:1 with MP-TLR4 for 15 mins, then sorted) were incubated at 100k cells in 200 uL in cell culture media (10% HIFBS in RPMI) and incubated at 37°C and 5% CO2 for varying time points. Then naïve BMDCs (1 million in 1 mL) were mixed with FRs or nFRs separated by a 1 um transwell insert or without for 16 hrs. Cell supernatants were tested via CBA for cytokine secretion and cells were tested via flow cytometry for CD80 (PerCP-Cy5.5 rat anti-mouse CD80, 1:100) and CD40 (APC rat anti-mouse CD40, 1:200), incubated for 1 hr at 4°C then washed 3X with PBS, then tested via flow cytometry.

*Adoptive Transfer of BMDCs* FRs and nFRs treated with MP-TLR4 similar to previous section were treated with DiL dye (1 ug/mL) and OVA (100 ug/mL) in 1 million cells in 1 mL cell culture media for 30 mins. Treated cells were then washed with PBS then concentrated at 1 million cells into 30 uL of HBSS and then immediately injected into C57BL/6 mouse footpads, one injection per footpad, two per mouse. 1 hr post injection, blood was taken for CBA analysis. 14 days later, mice were sacrificed, popliteal lymph nodes from both sides of the mouse were removed, disaggregated, stained and analyzed via Aurora spectral flow analysis.

*mRNA Whole Transcriptome Analysis*

For 16 h sequencing: BMDCs were incubated at a 1:1 ratio with MP^TLR-X^ (x = blank, 2, 4, 5, 7, 9, 2_4, 2_5,

2_7, 2_9, 4_5, 4_7, 4_9, 5_7, 5_9, 7_9) for 16 h. The FRs and nFRs were isolated via FACS. RNA was

extracted using a Direct-zol RNA-Microprep kit (Zymo), prepped using SMARTer® Stranded Total RNA-Seq

Kit v2 (Takara), and sequenced on a NextSeq550 (Illumina). RNA seq reads were mapped to GRCm38

mouse reference genome using STAR version 2.7.0b^6^ The resulting files from the alignment step above

were taken to evaluate transcriptional expression using subread::featureCounts with gencode transcript

annotation M19.^7^ The obtained count table was normalized and log fold change in expression was

generated using the edgeR package ^8^. Using the Cell Surface Protein Atlas’s database or mouse cell surface

protein, we identified proteins that were most frequently upregulated in the most MP^TLR-X^ dosing

conditions that met both the following criteria: 2-fold upregulation and pval <.05.^9^ For time series

sequencing: BMDCs were incubated at 1:1 ratio with MP^TLR-4^ for 15 min. The FRs and nFRs were isolated

via FACS, washed, and resuspended in media. At 0, 0.5, 1, 2, 4 h, the RNA was extracted using a Direct-

zol RNA-Microprep kit (Zymo). Sequencing was performed by the University of Chicago Genomics Core,

and BasePairTech’s DESEQ2 pipeline was used to align the reads to the mm10 genome, and compute the

differential expression and GSEA analysis. Using the Mouse Genomic Informatics database to acquire lists

of genes with 1) immune function 2) antigen presentation the immune response of the FRs and nFRs were

profiled by analyzing the differential expression, using only genes with 2-fold differential expression

change and pval < .05 for at least one of the timepoints.^10^

*Anti-OVA ELISA* Mouse anti-OVA IgG were measured using a commercially available kit from Alpha Diagnostic International according to the manufacturer’s instructions.

*Brefeldin A In vivo Experiment* 6 week, C57BL/6 female mice, 5 mice per experimental group were injected with liposomes containing brefeldin A or free Brefeldin A or PBS control. Mice were injected with 100 ug of Brefeldin A (either free or loaded equivalent) i.p. and then 1 hr later mice were injected i.p. with 10 ug R848 and 100 ug OVA in PBS. 14 days later this injection procedure was repeated. On day 21, mice were sac’d, blood tested for anti-OVA IgG and lymph and spleen analyzed.

*GpG Loaded Liposome In vivo Experiment* 6 week, C57BL/6 female mice, 5 mice per experimental group were injected with liposomes containing CpG or free CpG or PBS control. Mice were injected with 10 ug of CpG (either free or loaded equivalent) with 100 ug OVA i.p. On day 14, mice were sac’d, blood tested for anti-OVA IgG and lymph and spleen analyzed.

*R848 Loaded Liposome In vivo Experiment* 6 week, C57BL/6 female mice, 5 mice per experimental group were injected with liposomes containing R848 or free R848 or PBS control. Mice were injected with 10 ug of R848 (either free or loaded equivalent) with 100 ug OVA i.p. On day 14, this procedure was repeated. On day 21, mice were sac’d, blood tested for anti-OVA IgG and lymph and spleen analyzed.

*Intracellular Staining* Fresh suspensions of mouse spleenocytes were incubated with OVA peptides for 30 mins in ICS media (RPMI, 10% FBS, Penicillin/Streptomycin, 1x Non-Essential Amino Acids, 1 um B-mercaptoethanol, 1 mM HEPES and 1 mM Sodium Pyruvate. Then spleenocytes were incubated for 12 hrs with monensin (1 ug/mL), washed, fixed and stained for intracellular and extracellular antigens (CD4, CD8, INFγ and IL-4) according to the manufacture’s instruction for BD Cytofix/Cytoperm Kit.

*E.G7 OVA Tumor Model* C57BL/6 female mice were injected with 5x10^5^ E.G7 OVA expressing cells in PBS in the left flank. Mice were shaved to observe tumor growth. On day 7 and day 10, mice were injected with liposome formulation or free R848/OVA (10 ug/100 ug per mouse) i.p. Tumor volume was tracked via calipers 3 times per week and volume calculated by the formula Volume= (1/2) (length*(width)^2).^11^

*Statistics* Unless otherwise noted, all statistics were performed using a student’s t test with significance p<0.05. Poission distribution was used to calculate the standard distribution of discrete particles where Probability of Number of MP uptaken= P, κ= Number of particles uptaken per cell, λ= average number of MPs per cell for each experiment. $P=\frac{\lambda^{\kappa}e^{-\lambda}}{k!}$
